# Conserved Heterochromatin-like Structures with Local Regulators Mediate the Iron Stress Response in Mycobacteria

**DOI:** 10.1101/2025.08.23.671944

**Authors:** Alyssa M Ekdahl, Agata Turula, Jeremy W Schroeder, Rebecca Hurto, Quancheng Liu, Lydia Freddolino, Lydia M Contreras

**Affiliations:** Mcketta Department of Chemical Engineering, University of Texas at Austin, Austin TX; Department of Biological Chemistry and Department of Computational Medicine and Bioinformatics, University of Michigan Medical School, Ann Arbor MI

**Author notes:** To whom correspondence should be addressed. Correspondence may also be addressed to.

## Abstract

Recent studies have demonstrated the importance of dynamic heterochromatin-like regions in bacterial gene regulation, particularly for adaptation to changing environments. Here, we have measured the dynamic regulatory protein-DNA landscape of the tuberculosis vaccine strain, *M. bovis* BCG Pasteur, under the pathogenically-relevant condition of iron starvation. Our results capture for the first time the overall protein occupancy landscape of the genome of *M. bovis* BCG, identifying extended protein occupancy domains likely composed of diverse sets of nucleoid-associated proteins and transcription factors. Importantly, we find chromatin-directed regulation of stress-responsive genes like siderophores. Furthermore, through comparison with the free-living *M. smegmatis*, we identified a specific class of extended protein occupancy domains that are associated with conserved genomic regions across the two organisms, whereas regions with low protein occupancy often lack conservation. Our findings thus comprehensively reveal the contributions of both local regulators and chromatin structure to gene regulation and evolution in mycobacteria.

## INTRODUCTION

The mycobacterial genus contains numerous important pathogens, including *M. tuberculosis* and *M. bovis*, which make up the MTB complex^1^. *M. tuberculosis’* latent infections within human hosts are enabled by complex stress adaptation mechanisms^2,3^, hypothesized to contribute to its concerning rise in antibiotic resistance^4^.

Bacteria utilize multiplexed gene expression control for stress adaptation. Transcriptional regulation involves local and global protein binding events throughout the genome to direct or occlude binding of RNA polymerase (RNAP) to gene promoters^5^. Locally, transcription factors can bind near promoters (which we define as [-150, 70] bp surrounding transcription start sites in mycobacteria^6^) to control transcription of the neighboring genes through recognition of specific DNA sequences and structures^7^. Globally, nucleoid-associated proteins (NAPs) bind DNA with little sequence specificity to condense large regions into heterochromatin-like structures^8^. Mycobacteria have 5 major known NAPs: HupB and mIHF, which are among the five most prevalent proteins in the cell^8,9^; the xenogeneic silencer Lsr2^10,11^; the virulence regulator EspR^12^; and NapM^13^. NAP-directed stress responses have been previously observed in mycobacteria, where upon exposure to antibiotics, fluorescently labeled HupB and mIHF visibly changed binding patterns and chromatin structure in the model organism *M. smegmatis*^14^. Thus, transcription factors and NAPs play dynamic roles in coordinating stress responses in mycobacteria.

A large ChIP-seq study in *M. tuberculosis* documented genome-wide binding sites for 206 likely transcriptional regulators^6,15^. This study yielded the most comprehensive dataset available for *M. tuberculosis*, contributing alongside other individual ChIP-seq studies (such as for LexA^16^, mIHF^17^, and HupB^18^). While these studies have vastly expanded our understanding of *M. tuberculosis* transcriptional regulation, most ChIP-seq datasets have been collected under basal conditions, thus missing changes in the protein occupancy landscape in response to stress.

A recent method, *in vivo* protein occupancy display – high resolution (IPOD-HR), was developed to identify all DNA regions bound by proteins at a given time, agnostic to protein identity^19^. Since its development in *E. coli*, IPOD-HR has also been applied to the gram-negative pathogen *Vibrio cholerae*^20^ and the gram-positive model organism *Bacillus subtilis*^21^. By merging information from prior datasets, IPOD-HR has successfully mapped activities of known and novel transcription factor regulons regulating both protein-coding genes^19^ and stress-responsive sRNAs^22,23^. IPOD-HR has also captured NAP dynamics like large DNA regions densely bound by proteins, termed extended protein occupancy domains (EPODs)^8,21^. These EPOD regions contain high levels of bound protein covering genomic regions >1kb and are hypothesized to hinder and silence RNA polymerase (RNAP) function^8,20,21^. The prevalence of EPODs as captured by IPOD-HR across several bacterial species has supported the prevalence of heterochromatin-like structures for stress responsive gene regulation in bacteria.

In this work, we aimed to characterize the NAPs and transcription factors controlling stress responses in mycobacteria. We applied IPOD-HR alongside RNA-seq to the tuberculosis vaccine strain *M. bovis* BCG Pasteur and compared our results to existing ChIP-seq datasets. We found the NAPs HupB and mIHF are the primary contributors to most EPODs. Furthermore, we studied iron starvation and oxidative stress responses and found that many important processes, such as siderophore synthesis, are regulated through coordinated activities of transcription factors and dynamic EPODs. Lastly, by comparing to the free-living model organism, *M. smegmatis* mc^2^155, we found that heterochromatin-like structures in *M. bovis* BCG correlated with cross-species sequence conservation. In summary, this work greatly expands the knowledge of stress-responsive chromatin structures of *M. bovis* BCG and reveals the cross-species conservation of EPODs in mycobacteria.

## RESULTS

### IPOD-HR captures genome-wide DNA-protein interactions in Mycobacterium bovis BCG

To obtain information on global regulatory dynamics governing the transcriptome of mycobacteria, we performed *in vivo* Protein Occupancy Display – High Resolution, or IPOD-HR (which includes RNAP ChIP-seq), alongside RNA-seq as previously described^19^ during unstressed and stressed culture conditions. IPOD-HR provides a snapshot of the DNA-protein landscape across the genome *in vivo* alongside RNAP binding (**Figure 1A**). Briefly, cells are first treated with rifampicin to stall RNAP at promoters, then proteins are crosslinked to the DNA. DNA-protein complexes are isolated by phenol-chloroform phase separation where the interphase is extracted. Sequencing of the isolated DNA fragments, normalized by the total input DNA, produces a genome-wide snapshot of protein binding events along the DNA at the sampled time, however without identification of the proteins. With the addition of RNA-seq, the transcriptional effects of total protein and RNAP occupancies can be interpreted. Throughout this study, we refer to two types of protein occupancy datasets: IPOD, representing the total protein occupancy signal before RNAP subtraction, and IPOD-HR, representing the protein occupancy with RNAP contribution to the IPOD signal removed.

**Figure 1.**
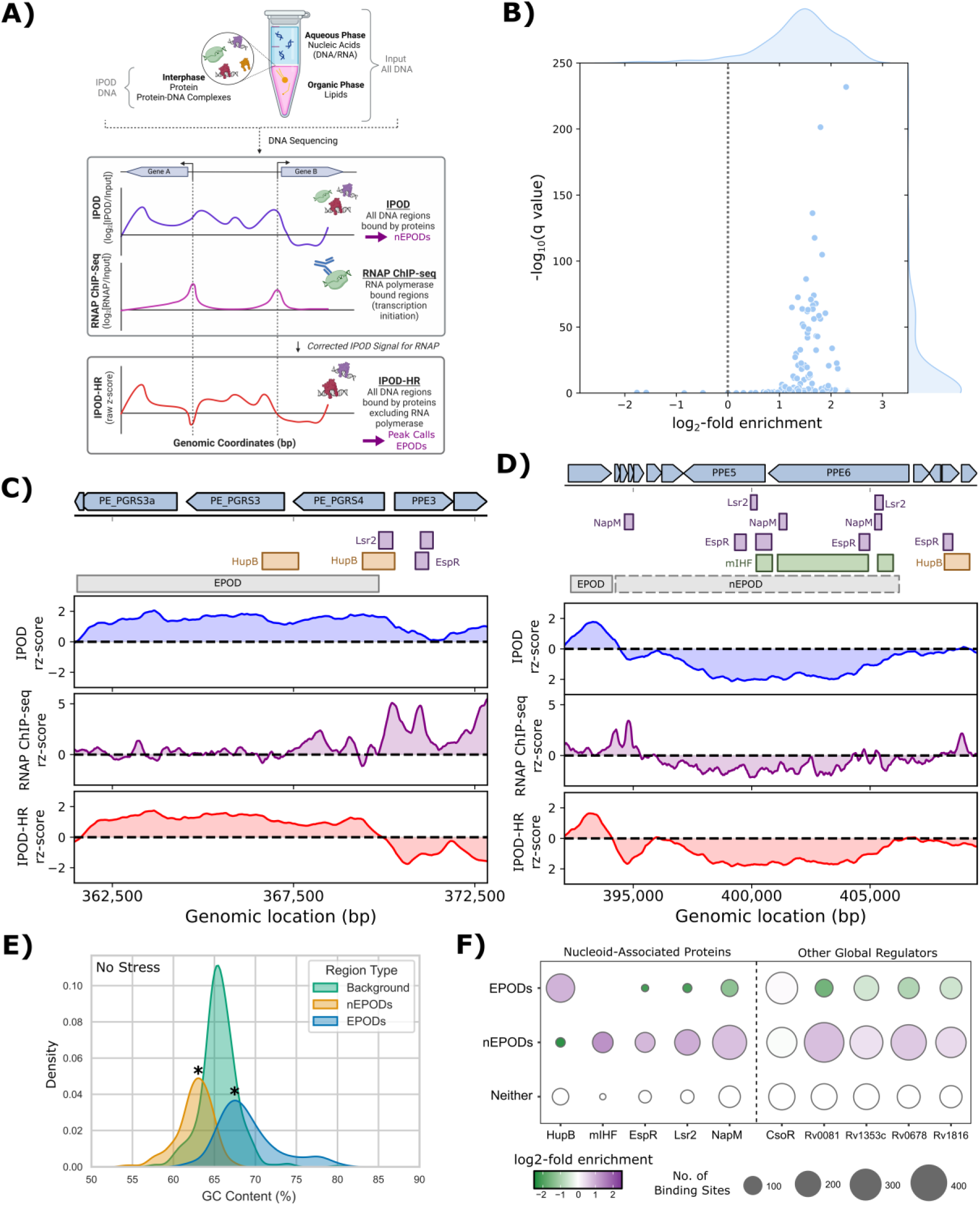
IPOD-HR captures global and local protein dynamics along *M. bovis* BCG genome. **A)** A simplified diagram of the IPOD-HR method, starting at the phase separation step (see Methods for details). The DNA sequenced from the interphase is normalized by the total DNA input via log_2_ ratio to achieve the IPOD protein occupancy signal (*i.e.*, a value of 0 means an equivalent fraction of that DNA region is in the interphase as is in the entire DNA in the sample, positive values mean enrichment at the interphase while negative values mean de-enrichment). The result is a snapshot of all protein binding events across the genome at a given time (IPOD, blue). RNAP ChIP-seq (purple) is used to adjust the IPOD signal to subtract RNAP contributions. The resulting IPOD-HR signal (red) is therefore representative of all non-RNAP protein binding events. Negative EPODs (nEPODs) are called from the IPOD signal, while peaks and positive EPODs are called from the IPOD-HR signal. Created in BioRender.com/4xxkryy. **B)** Comparison of peaks called from IPOD-HR occupancy to known ChIP-seq mapped binding sites, plotted for enrichment values (x-axis; log_2_-fold enrichment relative to the overall fraction of the genome covered be peaks) and statistical significance of the overlap (y-axis; based on a binomial test of the count of peaks overlapping known binding sites for each regulator, comparing the rate with the baseline rate of peak coverage; -log_10_ FDR-corrected p-values are shown). Out of 640 possible comparisons, across the 159 regulators and four conditions, 342 showed a significant correspondence at an FDR threshold of 0.1, and all but one of those significant cases showed an enrichment in overlap between binding sites and IPOD-HR peaks. **C-D)** Examples of positive (C) and negative (D) EPODs mapped during non-stress conditions. Nucleoid-associated protein (NAP) binding sites are indicated with colored bars (orange: HupB, green: mIHF, purple: all other NAPs). The grey horizontal bars indicate (n)EPOD regions. The IPOD (top, blue) and IPOD-HR (bottom, red) panels are plotted as 250 bp moving average, while RNAP ChIP-seq (middle, purple) plotted as 50 bp moving average. Note the relative lack of RNAP binding within the EPODs and nEPODs, consistent with transcriptional silencing. **E)** Density plot of the GC content within EPODs (blue) and nEPODs (orange) compared to the rest of the genome (background, green) in no stress conditions. Significance between EPODs and nEPODs versus background was performed by permutation tests (see Methods; *: p < 0.05). **F)** Dot plot of the 5 well established NAPs (left) and the 5 global regulators with highest number of ChIP-seq mapped binding sites (right). Common name or the *M. tuberculosis* homolog gene name listed for each. Dot size denotes the number of prior ChIP-seq annotated binding sites that overlap with loose EPODs, nEPODs, or neither (top to bottom). Dot color for the EPODs and nEPODs rows denotes the enrichment score (log_2_ ratio of regulator binding sites occurring within a (n)EPOD compared to random chance, positive values indicate enriched binding within (n)EPODs while zero or negative values indicate regulator binding sites are not enriched within (n)EPODs). Notice that HupB and mIHF show enrichment of binding in EPODs and nEPODs, respectively, while the remaining NAPs (EspR, Lsr2, and NapM) also show lesser nEPOD enrichment.

To assess the concordance of our IPOD-HR occupancy data with known transcription factor binding sites in *M. bovis* BCG, we cross-referenced the local peak calls obtained from IPOD-HR (**Table S1**) with a comprehensive compendium of previously obtained ChIP-seq data in *M. tuberculosis*^15–18,24^. As seen in **Figure 1B**, the vast majority of known regulators showed enrichment of overlaps with IPOD-HR binding sites. In agreement with what we have previously observed for other organisms such as *E. coli*^19^, local IPOD-HR binding peaks appear to correspond to transcription factor binding sites in many cases (**Figure S1A-C**). Comparing changes in protein occupancy around promoter regions across conditions is thus expected to provide important information on the dynamics of local regulator binding.

### The nucleoid-associated proteins (NAPs) HupB and mIHF are major components of two distinct types of extended protein occupancy domains (EPODs)

Extended protein occupancy domains (EPODs), in which large continuous segments of the genome (>1kb) display high protein occupancy, were observed in *M. bovis* BCG as previously found in other bacteria (**Table S1**). An example is shown in **Figure 1C**, in which an EPOD occupied an operon encoding PE-PGRS3 and PE-PGRS4. HupB, the HU homolog of *E. coli*, has two previously ChIP-seq mapped binding sites overlapping the EPOD region, likely contributing to the high protein occupancy. Similarly, negative EPODs (nEPODs) in which long segments of the DNA display low IPOD signal, were also observed throughout the genome. Negative EPODs arise due to large-scale binding of proteins that partition out of, rather than into, the phenol-chloroform interphase, resulting in negative IPOD signals. Thus, EPODs and nEPODs are hypothesized to function similarly, just composed of proteins with different physiochemical properties. An example of such a region is the operon encoding PPE5 and PPE6 (**Figure 1D**), which showed a prolonged region of negative IPOD signal coinciding with minimal RNAP occupancy. Numerous NAPs are known to bind in this nEPOD region, such as the mycobacteria integration host factor (mIHF) and the virulence regulator EspR.

As previously observed in *V. cholerae*^20^, the average GC content was significantly higher in EPODs and lower in nEPODs compared to the rest of the genome (**Figure 1E**). The difference in GC content between regions support different bound protein compositions based on DNA binding recognition sequences. The binding site distributions and enrichments in (n)EPODs for the 5 established NAPs and other global regulators are provided in **Figure 1F** (all regulators in **Figure S1D**, **Table S2**). As expected, the NAPs preferentially bound in EPODs or nEPODs while other global regulators bound throughout the genome without preference. Notably, HupB and mIHF were the most enriched NAPs contributing to EPODs and nEPODs, respectively, with the other NAPs predominantly contributing to nEPODs. In fact, HupB was the most EPOD enriched regulator studied, where 41% of EPODs in the nonstress condition overlapped at least one HupB binding site. In contrast, a striking 94% of nEPODs overlapped at least one mIHF binding site. Thus, for the remainder of the manuscript, we refer to nEPODs as mIHF-type nEPODs, while keeping the understanding that nEPODs may also contain additional proteins beyond mIHF.

### Iron stress response is regulated by dynamic changes in protein occupancy and composition

We next investigated stress-responsive changes to the DNA-protein landscape in *M. bovis* BCG. We first tested the oxidative stress response known in prior literature to cause large-scale DNA condensation in mycobacteria^25^ (discussed in **Supplementary Information**, **Figure S2** and **Figure S3**). As expected, EPOD and nEPOD regions that formed during stress (188 and 295 EPODs and nEPODs, respectively) far outnumbered those that dissipated during stress (120 and 67), likely to protect essential operons from ROS damage. We then investigated the iron starvation response, a common stress imposed by host cells to starve invaders of essential nutrients. Iron starvation is of particular interest as it has been implicated in triggering dormant-like growth of *M. tuberculosis*^26^, a behavior associated with long-term infections. For iron starvation stress (“iron stress” for brevity), a previously established iron starvation procedure was followed^26,27^. HupB has previously been associated with iron-dependent expression^28^, and thus we expected EPOD regions to change between iron-rich and poor conditions.

As anticipated, iron stress resulted in protein occupancy changes (**Figure 2A**) alongside the expected transcriptomic stress response (*e.g.*, enriched “cellular response to stress” term in **Figure 2B**, RNA-seq results in **Table S3** and **Figure S3D-F**, Gene Ontology analyses in **Table S4**). However, as opposed to DNA condensation observed in oxidative stress, we observed the opposite during iron stress. EPODs and nEPODs were statistically larger on average in the iron-rich (3.6 and 5.4kb) compared to the iron-poor condition (2.8 and 3.7kb, respectively), supporting the hypothesis of more open chromatin structures in the iron-starved condition.

**Figure 2.**
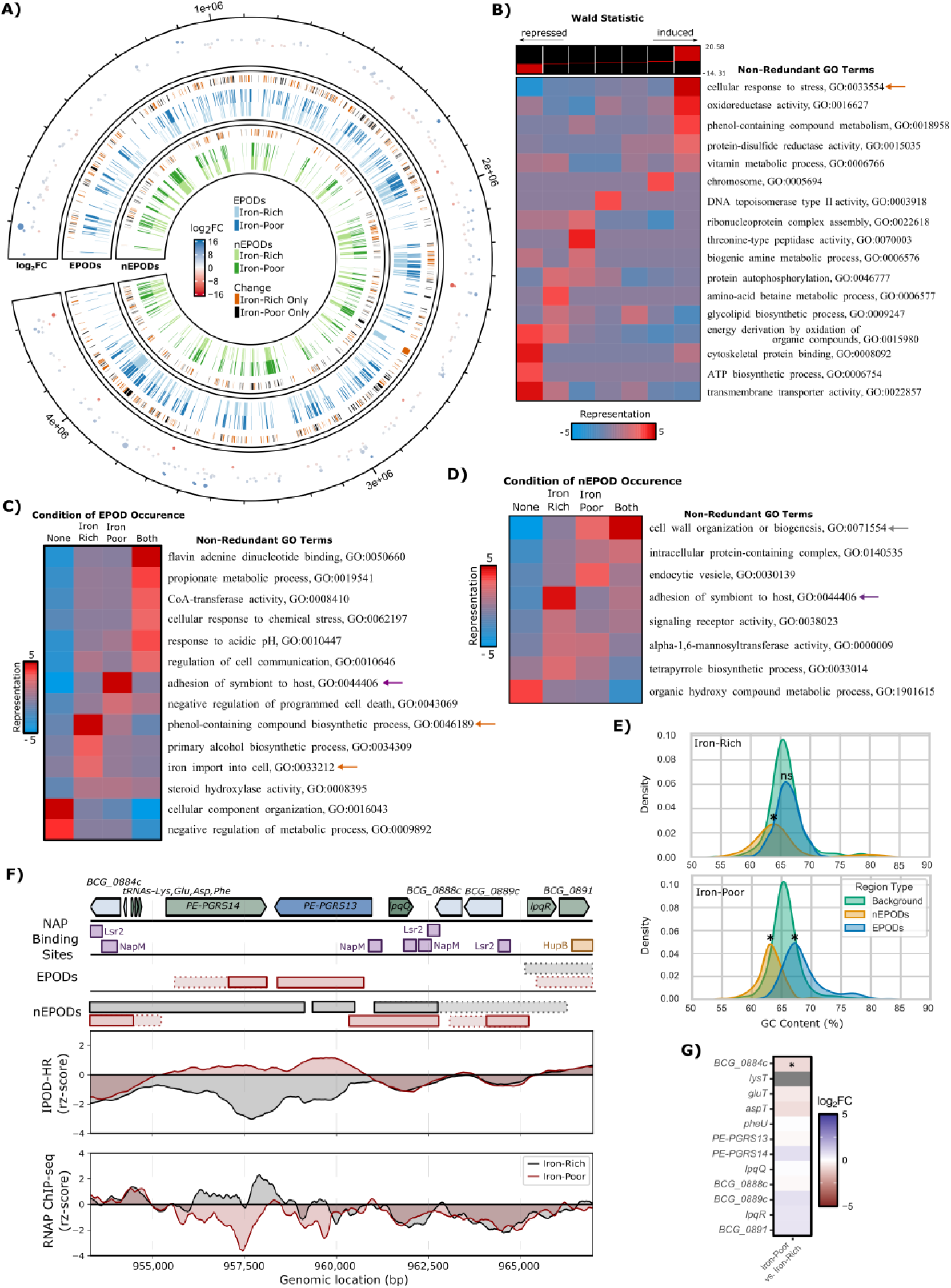
Extended protein occupancy domains show dynamic regulatory behavior alongside iron availability in *M. bovis* BCG. **A)** Genome-wide depiction of the significant transcriptional changes (outer, p-adj < 0.05, blue is induced, red is repressed), EPODs (dark blue = iron-poor, light blue = iron-rich), and nEPODs (dark green = iron-poor, light green = iron-rich). The outer edge of the EPODs and nEPODs rings indicate regions >1kb that change between conditions (orange = (n)EPOD region only called in the iron-rich condition, black = region only called in the iron-poor condition). **B)** Gene Ontology analysis of the RNA-seq differential expression results for iron stress relative to the corresponding non-stress (*i.e.*, iron-rich) condition. Noted by color coordinated arrows (with panels C-D) are cellular pathways discussed in the main text to be regulated by EPODs and nEPODs; the top of the panel shows the value ranges of the Wald statistic represented in each bin. **C-D)** Gene Ontology results for genes within (C) EPODs and (D) nEPODs in none, iron-rich only, iron-poor only, or both conditions (left to right columns). Enriched GO terms that passed the redundancy filter are provided with colors representing the enrichment score (red: overrepresented, blue: underrepresented). Cellular pathways of interest are annotated by arrows: EPOD-dependent regulation of iron homeostasis (orange) and consistent nEPOD regulation of cell wall composition (grey). The GO term, adhesion of symbiont to host, is highlighted with the purple arrow as a cellular process of interest as it is consistent with stress-responsive EPOD regulation (*i.e.*, genes within EPODs during iron stress but not in nonstress conditions). Notably, this term also appears in the nEPOD Gene Ontology analysis with the opposite behavior (within nEPODs during no stress conditions but not in iron stress) suggesting genes within this cellular pathway are switching from within nEPODs to EPODs during iron stress. **E)** Density plots of the GC content of EPODs (blue) and nEPODs (orange) compared to the rest of the genome (green) during iron-rich (top) and iron-poor (bottom) conditions. Significance of GC content for EPODs and nEPODs compared to background was determined by permutation test (p-value < 0.05 annotated with an asterisk). **F)** The protein occupancy results for an example operon, encoding PE-PGRS 13 and 14, that contributed to the enrichment of the adhesion of symbiont to host GO term in stress-responsive (n)EPODs. From top to bottom: prior ChIP-seq determined NAP binding sites (HupB: orange, others: purple), EPODs, nEPODs, IPOD-HR protein occupancy (rz-score), and RNA polymerase ChIP-seq (rz-score). EPODs and nEPODs are indicated by condition-colored boxes (black: iron-rich, red: iron-poor) with borders signifying achieved threshold (solid: strict, dashed: loose; see Methods for EPOD and nEPOD loose and strict threshold definitions). IPOD-HR and RNAP ChIP-seq results also colored by condition, as noted. Observe the dynamic EPODs and nEPODs for the genomic region encoding PE-PGRS13 and 14: contained within EPODs during iron-poor condition, and within nEPODs in iron-rich condition. In both cases, RNAP activity is muted. **F)** Resulting differential expression during iron stress associated with the PE-PGRS13 and 14 operon, (log_2_FC, asterisk to note p-adj < 0.05). Although the operon switches between nEPODs to EPODs during iron stress, the lack of expression change of PE-PGRS13 and 14 along with the lack of RNAP activity support the transcriptional silencing of this operon by EPODs and nEPODs.

We identified the cellular pathways enriched for by genes within EPODs (**Figure 2C**) and mIHF-type nEPODs (**Figure 2D**) between conditions. Genes within EPODs in only one of the compared conditions suggest dynamic regulation by NAPs binding or unbinding the DNA in response to iron starvation. As anticipated, enriched pathways associated with iron homeostasis such as iron import and siderophore synthesis (represented by the “phenol-containing compound biosynthesis” term) were within EPODs during iron-rich conditions and were in open chromatin during iron-poor conditions to allow expression. Alternatively, mIHF-type nEPODs appeared to regulate cell wall composition, with muted dynamics compared to EPODs.

Notably, the “adhesion of symbiont to host cell” GO term was enriched in both stress-responsive EPODs and nEPODs. Genes that made up this enriched pathway encode PE-PGRS proteins unique to mycobacteria. PE-PGRS proteins are presented on the cell exterior to interact with the host immune system^29^ and contain high GC-rich sequences, consistent with the high GC content tail apparent in EPODs only during iron-poor conditions (**Figure 2E**). As exemplified for the PE-PGRS13 and PE-PGRS14 operon (**Figure 2F**), protein occupancy and composition changed between conditions (*i.e.*, EPOD in iron-poor to nEPOD in iron-rich conditions). The changing protein composition between EPODs and mIHF-type nEPODs kept transcription silenced in both conditions (**Figure 2G**). This trend holds for numerous other PE-PGRS operons (**Figure S4A-C**). The differing protein occupancy dynamics of PE-PGRS loci may be attributed in part to WhiB4 binding, and potentially its interplay with mIHF^25^. In total, 376 genes switched between being contained within an EPOD to an nEPOD or vice versa during iron stress, including proteins involved in monooxygenase activity as well as aminoglycan biosynthesis (**Figure S4D**), suggesting the shifting protein composition may be a more widespread regulatory phenomenon beyond the PE family.

### Conservation of heterochromatin-like regions between free-living *M. smegmatis* and *M. bovis* BCG suggest evolutionary advantage

To determine if the stress-responsive EPOD regulation we observed in *M. bovis* BCG is ubiquitous in the mycobacteria family, we next studied the iron stress response in the free-living and fast-growing *Mycobacterium smegmatis,* in which its lack of host iron restriction yields an interesting contrast to the iron regulatory requirements of *M. bovis* BCG. Indeed, much of the iron stress response differed between the organisms, where only 43 out of the 2,573 total homologous genes (excluding duplicates, **Table S5**) were differentially expressed in both organisms (**Table S3**), including the expected iron importers (such as *lrtAB* and mycobactin) as well as components of the metal transporting ESX-3 Type VII secretion system (**Figure S5A-B**).

The protein compositions of (n)EPODs in *M. smegmatis* are likely similar, since HupB and mIHF are well-conserved between the organisms. Consistently, nEPODs remained AT-rich while EPODs remained GC-rich, but without the previously discussed dynamics of the high GC regions (likely since non-pathogenic mycobacteria lack PE-PGRS proteins, **Figure S5C**). Iron-dependent changes in (n)EPODs in *M. smegmatis* still supported some EPOD-dependent regulation, albeit to a lesser extent (**Figure S5D-F**).

One operon exemplifying EPOD-dependent regulation in response to iron starvation was the mycobactin synthesis operon, shown in **Figure 3A**. Specifically apparent in *M. bovis* BCG was the lack of EPODs in the iron-poor condition which coincided with high RNAP activity and gene expression throughout the transcribed operon (**Figure 3B**). Importantly, multiple known HupB binding sites have been previously identified to regulate siderophore transcription, alongside the iron-dependent IdeR transcription factor^28,30^. Alternatively, we observed different protein occupancy behavior in the homologous mycobactin operon in *M. smegmatis*: EPODs in *M. smegmatis* were located throughout the operon in iron-rich conditions (albeit with weaker occupancy than in *M. bovis*) yet retained high protein occupancy in the iron-poor condition (**Figure 3C**). The lack of EPOD-directed regulation of the mycobactin operon in *M. smegmatis*, however, did not coincide with transcriptional silencing, as gene expression did still increase in the iron-poor condition as expected in **Figure 3D**. Thus, the mycobactin operon in *M. smegmatis* is likely controlled predominantly by local transcription factors, such as IdeR, which has a known binding site between *mbtA* and *mbtB*. The other non-conserved siderophore operons between the organisms further support this observation (**Figure S6**). These siderophore examples demonstrate a potential difference in the regulatory use of EPODs between the two species: *M. bovis* BCG used large EPODs (>5kb) to silence transcription during iron-rich conditions that fully dissipated in iron-poor conditions, while *M. smegmatis* used short and less stress-responsive EPODs likely relying more on local regulators to toggle transcription.

**Figure 3.**
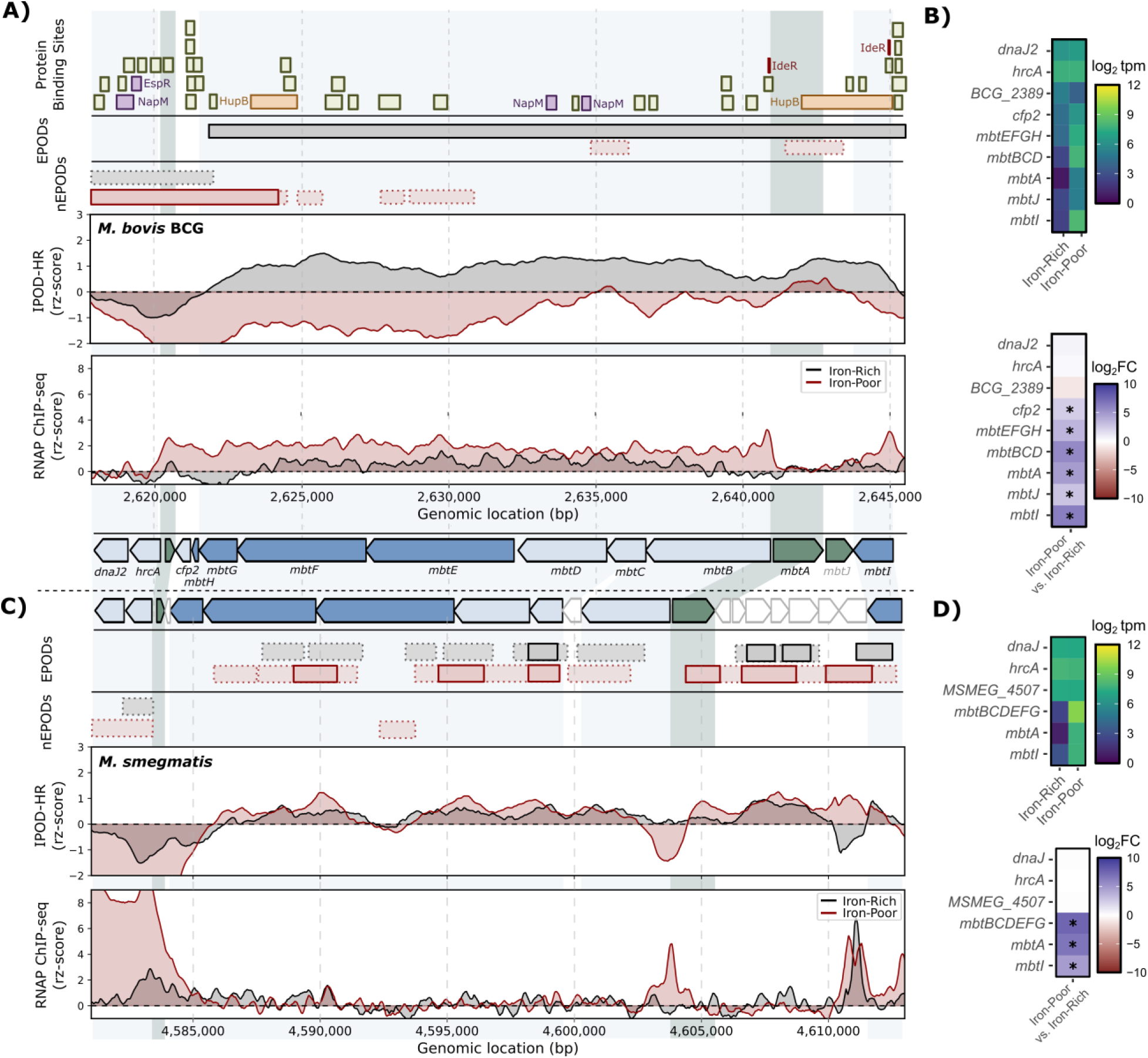
Siderophore biosynthesis operons regulated by iron-dependent EPODs in *M. bovis* BCG while less clear in *M. smegmatis*. **A)** The IPOD-HR results of the siderophore operon encoding mycobactin in *M. bovis* BCG. Information provided from top to bottom: prior ChIP-seq determined protein binding sites, EPODs, nEPODs, IPOD-HR protein occupancy (rz-score), and RNA polymerase ChIP-seq (rz-score). Binding sites for nucleoid-associated proteins (HupB: orange, others: purple), the iron-dependent repressor IdeR (maroon), and other proteins (olive green) are indicated. EPODs and nEPODs are indicated by condition-colored boxes (black: iron-rich, and red: iron-poor) with borders signifying achieved threshold (solid: strict, dashed: loose). IPOD-HR and RNAP ChIP-seq results also colored by condition, as noted. The encoded gene positions noted at the bottom of the figure with homologous genes in *M. smegmatis* are shaded green and blue. **B)** Mycobactin operon gene expression as log_2_(transcripts per million, tpm) for iron-rich and iron-poor conditions (shown on top), as well as the corresponding log_2_(fold change) in expression under iron stress (bottom) with significance (p-adj < 0.05) annotated with stars. **C)** The homologous operon for mycobactin synthesis in *M. smegmatis* with accompanying results with same organization as panel A for iron-rich (black) and iron-poor (red) conditions. Non-conserved regions in *M. smegmatis* are not shaded. **D)** Gene expression in *M. smegmatis* for mycobactin operon, organized as in panel B.

Indeed, we found that genome-wide EPOD and nEPOD maps (**Figure 4A**) for *M. bovis* BCG and *M. smegmatis* highlighted the global differences between organisms (**Figure S7**, **Table S5**). With respect to the presence of (n)EPODS, patterns begin to emerge, such as the denser appearance of the EPOD and nEPOD regions in *M. bovis* BCG compared to *M. smegmatis.* Indeed, *M. bovis* BCG had statistically larger EPOD and nEPOD regions in iron-rich conditions, and nEPOD regions in iron-poor conditions (**Figure 4B**); in iron-rich conditions, the average EPOD length in *M. bovis* BCG was 3.6kB, a ∼70% increase compared to 2.1kb in *M. smegmatis*.

**Figure 4.**
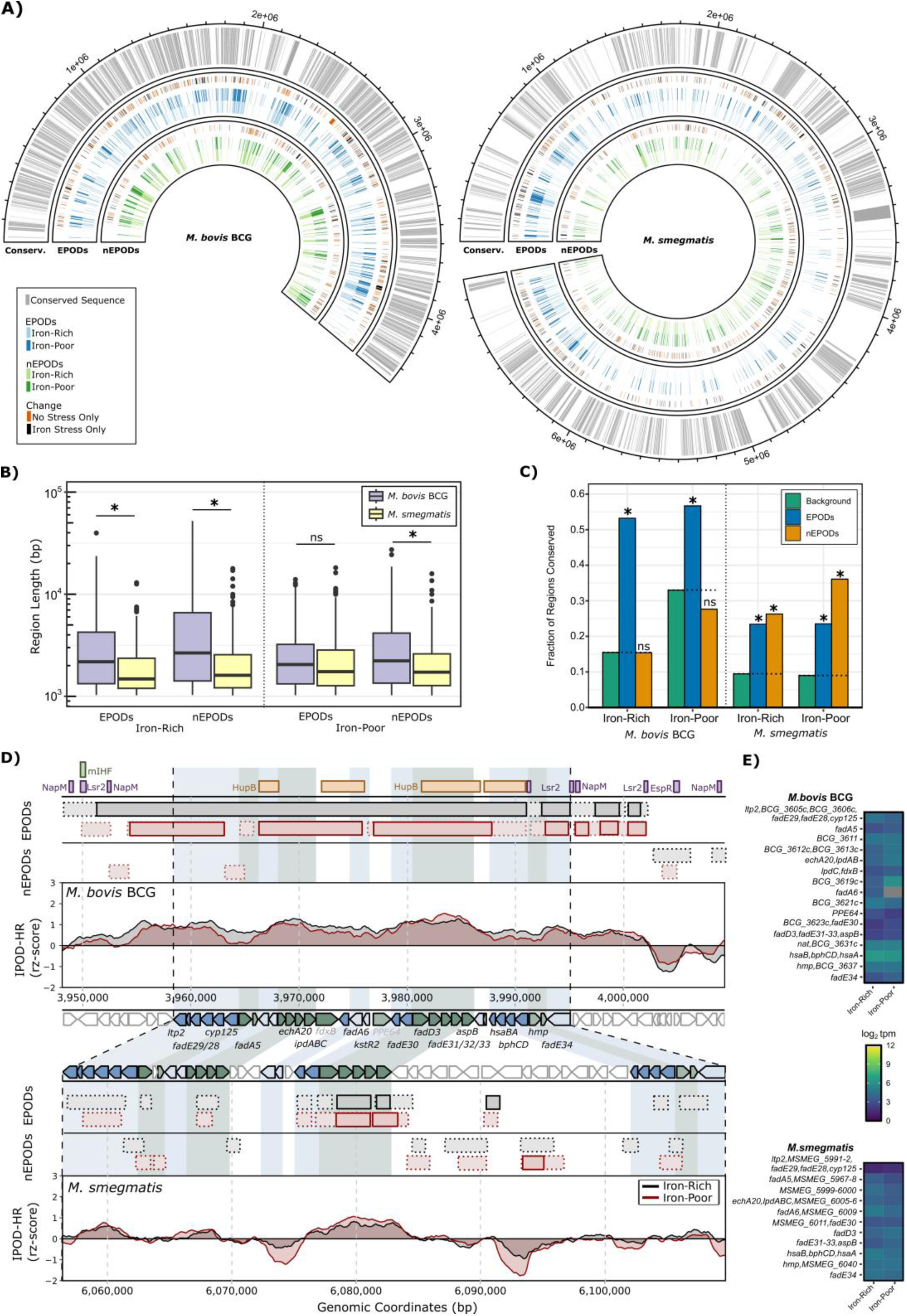
Heterochromatin-like structures occur in sequence conserved regions of *M. bovis* BCG and *M. smegmatis*. **A)** Genome-wide depiction for *M. bovis* BCG (left) and *M. smegmatis* (right) of the sequence conservation between organisms (outer, grey), EPODs (middle, light blue = iron-rich, dark blue = iron-poor), and nEPODs (center, light green = iron-rich, dark green = iron-poor). Changes in (n)EPOD regions are depicted on the outer portion (orange = (n)EPOD region present during iron-rich condition only, black = present during iron-poor condition only). **B)** The length distributions of EPODs and nEPODs in *M. bovis* BCG (purple) and *M. smegmatis (*yellow) in both iron-rich (left) and iron-poor (right) conditions. Statistical significance between organisms by Welch’s t-test are illustrated with an asterisk for p-values < 0.05. **C)** The fraction of EPODs and nEPODs with >50% sequence conservation between organisms as compared to background regions (limited to regions within the range of (n)EPOD lengths for the given condition). Significance from background by Fisher Exact Test indicated with an asterisk for p-values < 0.05. **D)** The IPOD-HR protein occupancy (rz-scores) for the operon encoding for various lipid metabolism genes in *M. bovis* BCG (top) and *M. smegmatis* (bottom) in iron-rich (black) and iron-poor (red) conditions. Homologous genes shown by shading (green: forward strand, and blue: reverse strand) between organisms. EPODs and nEPODs plotted above each by colored horizontal bars. Known binding sites of nucleoid-associated proteins (NAPs) by ChIP-seq plotted at the very top of the plot (HupB in orange, mIHF in green, rest in purple). **E)** Gene expression (by log_2_tpm) of the cholesterol operon in panel D for *M. bovis* BCG (top) and *M. smegmatis* (bottom). Note *fadA6* had no detected transcripts in the iron-poor condition (grey).

We also observed a correlation between sequence conservation between organisms and (n)EPOD occurrence. EPODs in *M. bovis* BCG and both EPODs and mIHF-type nEPODs in *M. smegmatis* were enriched in conserved DNA regions between the organisms (**Figure 4C**, **Table S5**). In fact, EPODs had odds ratios of 3.4 and 1.7 in iron-rich and iron-poor conditions, respectively, to occur in highly conserved regions of the genome in *M. bovis* BCG compared to background (notably, nEPODs lacked significant correlation with conservation). Similarly, EPODs and nEPODs in *M. smegmatis* were also more likely to occur in conserved regions in both conditions (odds ratios between 2.5 - 4.1 compared to background). Literature speculates that the MTB complex organisms evolved from a non-pathogenic fast-growing mycobacteria species^31^, such as *M. smegmatis*. As is common for pathogens, *M. bovis* BCG has a much more condensed genome than *M. smegmatis*, hypothesized to be due to selective pressures in human hosts to optimize genome size^32,33^. Thus, EPOD-regulated regions were more likely to be conserved through the genome reduction in *M. bovis* BCG, either reflecting their biological importance or a mechanistic link between high protein occupancy and reduced gene loss.

A striking example of the conservation patterns of EPODs occurred in the operon encoding lipid metabolism genes^34^ (**Figure 4D**, with another cross-species example in **Figure S8**). *M. bovis* BCG had static high protein occupancy in all tested conditions over a region >35kb, while *M. smegmatis* had EPODs covering conserved regions with *M. bovis* BCG and unbound DNA regions in non-conserved regions, resulting in shorter dispersed (n)EPODs. Although *M. smegmatis* contained open chromatin in which RNAP was able to bind and transcribe genes, the conserved regions sustained high protein occupancy in both organisms to silence transcription (**Figure 4E**). Thus, the conserved NAPs, HupB and Lsr2, with binding sites overlapping the homologous genes, likely bound and suppressed transcription in *M. smegmatis,* albeit not showing the uniform occupancy seen in *M. bovis* BCG. The enrichment of the EPODs in conserved genomic regions between *M. smegmatis* and *M. bovis* BCG suggests at least two exciting possibilities that we cannot distinguish: (1) there is some direct protection of these regions during genomic collapse of the *M. tuberculosis* complex organisms (*e.g.*, resistance to loss via recombination), or (2) genes contained within EPODs encode proteins of conserved biological need. Also noteworthy is the lack of correlation between sequence conservation and nEPODs in *M. bovis* BCG in both conditions; another feature to consider in future investigations as to how chromatin structure by (n)EPODs could impact the location and frequency of novel DNA integration.

## DISCUSSION

This study captured the genome-wide protein landscape regulating stress responses in the tuberculosis vaccine strain *M. bovis* BCG Pasteur. As previously demonstrated in other organisms, coupling IPOD-HR with previously mapped binding sites by ChIP-seq can inform protein identities making up the IPOD-HR signal^19–21,23^ and identify orphan peaks without a mapped binding site suggestive of uncharacterized DNA-binding protein activity. While informative, leveraging these two complementary datasets had some notable caveats worth considering when interpreting the data, discussed in **Supplementary Information**. Nevertheless, the IPOD-HR approach dramatically reduces the number of sequencing experiments needed to capture protein-directed stress responses while revealing coordinated, large-scale protein occupancy regions that are difficult to appreciate in single-protein ChIP experiments.

Across test conditions in our *M. bovis* BCG experiments, we observed a strong correspondence between local regulator occupancy peaks and known binding sites. IPOD-HR also identified extended protein occupancy domains (EPODs and nEPODs) made up of dense protein binding events across conditions. These regions are suggestive of heterochromatin-like DNA structures where nucleoid-associated proteins, or NAPs, compact large segments of DNA^8,9^. HupB was the only NAP found to be enriched for binding within EPODs, while mIHF (and to a lesser extent NapM, Lsr2, and EspR) was strongly enriched in nEPODs. Previous studies in mycobacteria have found sequence preferences for AT-rich regions (mIHF^17^, HupB^28^, Lsr2^35^, EspR^12^, NapM^13^) and GC-rich regions (WhiB4^25^). Since the GC content differed between EPODs and nEPODs, these regions were likely made up of differing protein compositions with different DNA binding recognition sequences.

We hypothesize that positive EPODs are frequently comprised of HupB while nEPOD are comprised predominantly of mIHF alongside the other identified NAPs. As previously interpreted, EPODs typically silence transcription of the encompassed genes^8,21^; this functionality of EPODs generally holds in mycobacteria during stress (**Figure S3B**). Importantly, dynamic (n)EPOD regions were found to encompass genes of stress-importance (*i.e.*, relating to iron homeostasis and oxidative stress response) and virulence (*i.e.,* adhesion of symbiont to host). These dynamic regions are of particular interest as they suggest EPOD-dependent transcriptional regulation. In mycobacteria, some NAPs have been associated with the studied stresses, such as HupB for iron stress^28,30^ and Lsr2 for oxidative stress^36^. Yet, binding sites for these NAPs do not account for all of the dynamic regions, suggesting the involvement of other regulators or un-annotated binding sites for the known NAPs. NAPs typically have much wider binding sites than reported due to their broad, low-level binding not easily recognized by classical ChIP-seq analyses. There may also exist NAPs in mycobacteria which are either undiscovered^8^ or misclassified, perhaps like WhiB4 for the PE-PGRs operons^25^. Additionally, NAPs likely have changing binding sites during different stress conditions, as seen in prior studies^14,25^. Therefore, our results support contributions of stress-responsive proteins to the heterochromatin-like regulation yet to be characterized.

When compared to the free-living *M. smegmatis*, we found that many of our observations in *M. bovis* BCG were not ubiquitous across the mycobacterial genus. Most notably, we observed reduced reliance of *M. smegmatis* on stress-responsive EPOD regulation. We also found global sequence conservation of heterochromatin-like regions between the distant relatives. Others have hypothesized that the MTB complex organisms evolved from a non-pathogenic mycobacterium^31^, like *M. smegmatis*, in which the genome reduced in size by mammalian host selective pressures. It has been postulated that nucleoid structure could bias the evolutionary conservation and DNA integration likelihood of genomic regions^21,37^. We also speculate that the stronger reliance of *M. bovis* BCG on EPOD-mediated regulation may arise due to the relatively low entropy of the pathogen’s encountered environments (*i.e.*, limited combination of stresses in a mammalian host) compared to the less predictable free-living environment of *M. smegmatis*.

In conclusion, this study revealed the mechanistically complex protein-directed transcription regulation during stress in *M. bovis* BCG, requiring coordination of transcription factors alongside dynamic heterochromatin-like structures. Importantly, the siderophore and PE-PGRS operons emphasize remaining knowledge gaps involving dynamic nucleoid structures regulating virulence (and their protein-level compositions). Future studies are warranted to explore whether the unique heterochromatin-like regulation is conserved in the human pathogen, *M. tuberculosis*, and if its mechanisms can be specifically targeted by future therapeutics.

## ONLINE METHODS

### Cell Media

Normal liquid culture used Middlebrook 7H9 media supplemented with OADC and Tween-80 (4.7g Middlebrook 7H9 broth base, 2mL glycerol, 12g agar, 900ml DI water, 0.5mL Tween-80, 100mL OADC supplement [Sigma Aldrich] aseptically added after autoclave sterilization), with solid agar plates containing in addition 12g/L agar. All cell cultures grown at 37°C, with continuous shaking for liquid cultures. Cell pelleting performed at 3000rpm for 10-15 minutes.

For iron stress samples, Minimal Mycobacteria Media (100mL ADN Supplement [100mL DI water, 0.81g NaCl, 5g Bovine Albumin Serum Fraction V, 2g dextrose], 900mL DI water, 5g asparagine, 5g potassium phosphate monobasic, 25.2g glycerol, 0.545g Tween 80, pH adjusted to 6.8) was made following previous literature^26,27^. Briefly, metals were chelated from media overnight at 4°C using 50g/L Chelex-100 (Bio-rad) under moderate agitation. Vacuum filtration removed solid chelator. Media was then supplemented with metals (40g/L MgSO_4_, 0.1g/L MnSO_4_, 0.5g/L ZnCl_2_). Iron-rich media was also supplemented with 50μM FeCl_3_ for *M. bovis* BCG and 100μM FeCl_3_ for *M. smegmatis*. Since iron can leach from glass, all iron stress experiments performed in polypropylene tubes and flasks.

For oxidative stress samples, Middlebrook 7H9 media supplemented with 10% vol/vol ADN Supplement and 0.5mL/L tyloxopal was used. Tert-butyl hydroperoxide (TbHP) at 1mM concentration was used as the oxidizing agent as previously described^27^.

### *M. smegmatis* Iron Stress Cell Culture

*M. smegmatis* MC^2^ 155 strain (ATCC #700084) was streaked from frozen glycerol stocks onto 7H9 supplemented with OADC agar plates. Plates were incubated for 2-3 days. Single colonies (biological duplicates) were inoculated in Middlebrook 7H9 liquid media with OADC and Tween-80 and grown to saturation (2 days). Overnight cultures were then washed and resuspended in an equal volume of iron-poor minimal media. Cultures were seeded 1:100 into fresh iron-poor or iron-rich media to an OD_600_ ∼0.5 (12 hours after seeding), then sampled.

### *M. bovis* BCG Oxidative Stress Cell Culture

*Mycobacterium tuberculosis* variant *bovis BCG* strain TMC 1011 (ATCC #35734) was streaked from frozen glycerol stocks on 7H9 supplemented with OADC agar plates. Plates were incubated for 2-3 weeks. Single colonies (biological duplicates) were inoculated in Middlebrook 7H9 liquid media with OADC and Tween-80 and grown to saturation. Cultures were seeded 1:10 in fresh media and grown to late logarithmic phase (OD_600_ ∼1-2). Cultures were split and diluted to OD_600_ ∼0.6. Split cultures were either exposed to 1mM TbHP or no stress for 4 hours, then sampled.

### *M. bovis* BCG Iron Stress Cell Culture

The same saturated cell cultures from the oxidative stress cell cultures were used (biological duplicates), where cells were washed in equal volume iron-poor media. Following prior mycobacteria iron starvation protocols^26,27^, cultures were seeded 1:10 in fresh iron-poor media and grown to mid logarithmic growth (OD_600_ ∼0.6). Cultures were washed in equal volume iron-poor media. Cultures were split, seeded 1:10 in fresh iron-poor or iron-rich media and grown to OD_600_ ∼0.2-0.4. Deferoxamine mesylate (50μg/mL) was then added to the iron-poor samples to further chelate any remaining iron and grown for 24 hours, then sampled. Iron-poor samples experienced stunted growth as previously observed^26^, reaching OD_600_ ∼0.3, while iron-rich samples reached OD_600_ ∼0.7-1.0.

### Sample Preparation for RNA-Seq

Samples (5mL liquid cultures, 10mL culture for iron-poor BCG samples due to low ODs) were suspended in 2X volume RNAProtect (Qiagen) and incubated for 5 minutes at room temperature. Cells were then pelleted, supernatant poured off, and flash frozen for -80°C storage.

To extract RNA, cell pellets were resuspended in 1mL Trizol Reagent (Bio-Rad) and incubated with glass beads (250mg, 0.1 mm diameter) in screw-capped vials. Samples underwent bead beating for 2 minutes, then were chilled on ice. 24:1 Chloroform-isoamyl (300μl) was added to separate nucleic acids into top aqueous phase by microcentrifugation. Nucleic acids were precipitated overnight in isopropyl alcohol with Glycoblue (Ambion), then underwent ethanol washes (75% then 95%) prior to resuspension in nuclease-free water. Finally, DNAse I treatment was followed by an RNA column clean-up to remove genomic DNA contamination (Monarch RNA Clean-up kit). Quality checks of total RNA was performed by Nanodrop, then Bioanalyzer and Qubit fluorometry at the Genomic Sequencing and Analysis Facility (University of Texas at Austin).

### RNA Library Preparation, Sequencing and Analyses

Total RNA (250 ng each, 2 biological replicates per strain and condition) was depleted of ribosomal RNA using NEBNext rRNA Depletion Kit for Bacteria (E7850, NEB) per manufacturer’s protocol. The sequencing libraries were prepared per manufacturer’s protocol using NEBNext Ultra II Directional RNA Library Prep Kit for Illumina (E7760, NEB) except NEBNext Multiplex Oligos for Illumina (Unique Dual Index UMI Adaptors DNA Set 1; E7395L, NEB), diluted 1 to 100, were used as the adapters and Mag-Bind TotalPure NGS beads (M1378, Omega Bio-tek) were used for DNA clean-up.

RNA-seq datasets were quality checked by FastQC, then input into Rockhopper^38–40^ for alignment and *de novo* transcript assembly for each test condition separately. The transcriptomes generated from each condition were merged as follows: we collected all annotated genes from the existing GenBank annotations for the reference genomes noted above, plus all transcripts identified under any of the conditions using Rockhopper; any assembled transcripts with the same strandedness and more than 80% overlap were then pruned to avoid redundancy. The newly generated reference transcriptome thus includes some newly identified transcripts whose coordinates do not align with known gene boundaries. Characterization of these novel transcripts is outside the scope of the current study (annotated as new_##### with coordinates provided in **Table S3**). Having established a comprehensive reference transcriptome for each strain of interest, we then quantified transcript abundance on the newly generated transcriptome using kallisto version 0.48.0^41^, with a kmer size of 19, and 250 bootstrap samples. The resulting expression counts were input into sleuth ver. 0.30.1^41^ for differential expression calling via Wald tests. Significant differential expression of transcripts was defined as having a p-adj < 0.05 and |log_2_FC| > 1 (results provided in **Table S3**).

### Sample Preparation for IPOD-HR

As previously described in detail^19^, cell cultures were treated with 150 μg/mL rifampin for 10 minutes at 37°C with continuous shaking. Formaldehyde to 1% final concentration was added, incubated at room temperature for 30 minutes shaking (extended from the original protocol recommendation of 10 minutes). Excess glycine was then added to quench the crosslinking reaction (final concentration 0.333M) for 5 minutes at room temperature, shaking. Cultures were then chilled on ice and washed twice using phosphate-buffered saline. Final cross-linked cell pellets were stored at -80°C.

### Cell lysis and DNA preparation

Detailed instructions for IPOD-HR interface extraction, RNA polymerase chromatin immunoprecipitation, and crosslinking reversal and recovery of DNA have been previously published^19^, with slight modifications described here. Briefly, cross-linked pellets were resuspended in 750 μL lysis buffer [10 mM Tris pH 8.0, 1x cOmplete™, EDTA-free Protease Inhibitor Cocktail (Roche), 50 mM NaCl], and then transferred to 2mL Lysing Matrix B tubes (MP Biomedicals). Each tube was subjected to 3 rounds of bead beating (20sec, 3min on ice) in a Mini Bead Beater (Biospec products, Bartlesville, OK) at maximum speed.

Tubes were centrifuged briefly, and liquid was transferred to a new tube for RNaseA and DNase I (Fisher) treatment. For *M. smegmatis:* Each sample was brought to 600 μL using lysis buffer. Duplicate samples were then combined. To each sample, 10.8 μL of 100 mM MnCl_2_, 9.0 μL 100 mM CaCl_2_, 12 μL of RNase A (10 mg/mL) and 12 μL of DNase I (Fisher) was added and mixed. Samples were incubated on ice for 45 minutes before 100 μL of 500 mM EDTA was mixed in to stop DNase I digestion. Samples were clarified by centrifugation (16k r.c.f., 10 minutes, 4°C), then liquid was split into new microfuge tubes as follows: 50 μL for input, 500 μL for IPOD, and remainder (∼750 μL) for RNAP ChIP. For *M. bovis* BCG: 200 µL of additional lysis buffer was added to the beads, briefly vortexed and then centrifuged. The additional liquid was added to the samples. Samples were spun 30sec at 16K to pellet any insoluble material, and the liquid portion transferred to new tubes. The A_260_ (nanodrop) and total volume of each sample was determined. Per 600 μL of sample, 5.4 μL of 100 mM MnCl_2_, and 4.5 μL 100 mM CaCl_2_ was added and mixed. Then 6 μL of RNase A (10 mg/mL) per sample and 6 μL of DNase I (Fisher) per 9.9 OD_260_*mL was added per sample. Samples were incubated on ice for 30 minutes before 50 μL of 500 mM EDTA was mixed in to stop DNase I digestion. Samples were clarified by centrifugation (16k r.c.f.,10 minutes, 4°C). Samples were split so that equal amounts (OD_260_*mL) of stress versus non-stressed material per biological replicate were used for IPOD and RNAP ChIP, with 5-10% of the total material being used as input. Each sample for RNAP ChIP and IPOD were brought up to 500 μL using lysis buffer.

To each input sample, 450 μL of ChIP elution buffer (50 mM Tris pH 8.0, 10mM EDTA pH 8.0, 1% SDS) was added and mixed in. Inputs were maintained on ice until being placed at 65°C for 12 hours to reverse cross-links. To each RNAP ChIP sample, an equal volume of 2X IP buffer (200 nM Tris pH 8.0, 600 mM NaCl, 4% Triton X100) with 2x cOmplete™, EDTA-free Protease Inhibitor Cocktail and 10 μL of purified anti-*E. coli* RNA Polymerase β Antibody (1μg/μL, BioLegend) was added. ChIP samples were incubated overnight on a nutating platform at 4°C.

IPOD-HR interface extraction proceeded as described previously^19^, starting at the point of addition of 1 volume of 100mM Tris base and 2 volumes of 25:24:1 phenol:chloroform:isoamyl alcohol. RNAP ChIP samples were obtained as previously described^19^, except that after the antibody incubation overnight, 50 μL of pre-washed [once with 1000 μL 1X IP buffer + 0.1 mg/mL BSA (add buffer, mix by inversion, clear using magnetic stand, remove liquid)] protein G magnetic beads (NEB) were added to each sample instead of the specified protein G dynabeads. Reversal of crosslinks, recovery and quantification of DNA was performed as described^19^.

### DNA Library Preparation, Sequencing and Analyses

DNA obtained as input (10 ng), from IPOD (10 ng), or RNAP immunoprecipitation (1 ng for *M. smegmatis* and *M. bovis* BCG +/-iron, 2 ng for *M. bovis* +/- oxidative stress) was brought to 50 μL volume for each sample. Reagents from NEBNext Ultra II DNA Library Prep Kit for Illumina (NEB) and NEBNext Multiplex Oligos for Illumina Unique Dual Index UMI Adaptors DNA Set 1 (NEB) were utilized during library preparation. Samples prepared per manufacturer’s protocol for NEBNext Ultra II DNA Library Prep Kit using Multiplex Oligos for Illumina Unique Dual Index UMI Adaptors DNA Set 1 except for the bead mediated clean-up steps. The post-ligation clean-up step utilized 174 μL of either AxyPrep Mag PCR beads (Axygen Biosciences) or Mag-Bind TotalPure NGS beads (Omega Bio-tek) and 68 μL isopropanol added per sample (in microfuge tubes) for the sample to bead binding step in order to maintain the smaller fragments expected to be obtained from the IPOD protocol. Post amplification clean-up utilized 45 μL of AxyPrep Mag PCR beads (Axygen Biosciences) or Mag-Bind TotalPure NGS beads (Omega Bio-tek) for the sample to bead binding step. Sequencing libraries were eluted in 33 μL of 0.1X TE. QuantiFluor dsDNA dye (Promega) was used to determine DNA concentration while agarose gel electrophoresis was used to determine sample size distribution. The IPOD and RNAP ChIP DNA sequencing and analysis were performed as previously described and validated^19^. All reads were aligned to the *M. tuberculosis* variant *bovis* BCG str. Pasteur 1173P2 genome (Genbank Accession: AM408590.1) or the *M. smegmatis* mc^2^155 genome (Genbank Accession: CP000480.1).

### Feature Calling from IPOD and IPOD-HR Sequencing

Raw z-scores were averaged between the biological duplicates for each genomic coordinate at 5bp increments. Peak calling was performed on the IPOD-HR mean values using increasing thresholds at a peak width of 60 bp using the Python scipy.signal.peak_calling_cwt() method as previously benchmarked^19^. The peak threshold was selected to be 1 for all *M. bovis* BCG IPOD-HR samples based on two methods: 1) maximum K-L divergence across peak thresholds, and 2) precision-recall curves for all proteins (based on curated binding sites, see *Curation of Known DNA-Protein Binding Sites* section below) across peak thresholds 0 to 6 for the nonstress condition. The threshold of 1 was selected to optimize both the K-L divergence value as well as precision to recall over all proteins (peak calls in **Table S1**).

Extended Protein Occupancy Domain (EPOD) calling was performed as previously described^19^. Simply, 512 bp and 256 bp rolling window means were calculated for IPOD-HR rz-scores across the genome for each condition. From the 256 bp means, the 75^th^ and 90^th^ percentiles were defined as the loose and strict thresholds, respectively. Regions equal or greater than 1,024 bp long with median IPOD-HR rz-scores above the defined thresholds were considered a loose or strict EPOD (sliding window of 1,024 bp initially calculated, then expanded along the genome until the median occupancy fell below the respective loose or strict threshold). Similarly, the same was performed for negative EPOD (nEPOD) calling, however using the inverted IPOD (not IPOD-HR) signal, as the RNAP adjustment used to calculate the IPOD-HR signal can arbitrarily generate negative protein occupancy regions due to RNA polymerase signal subtraction. EPOD and nEPOD coordinates are given in **Table S1**.

### GC Content of EPODs and nEPODs

Permutation tests were performed to determine statistical significance between the populations of average GC content in EPODs and nEPODs, performed separately, compared to the rest of the genome. As performed previously^20^, we obtained 1000 randomized sets of genomic regions of same length and number as our real EPODs (or nEPODs) sets using bedtools shuffle^42^, excluding regions contained in the opposite nEPODs (or EPODs) sets. These randomized sets, or permutations, make up the null distribution of EPODs (or nEPODs). For each permutation, we calculated the difference in mean GC content compared to the respective background. We then compared the permutation results to the real EPODs (and nEPODs) GC content compared to background. The resulting P-value was calculated as the ratio of 1 plus the number of permutations with GC content at least as extreme as the real EPODs (or nEPODs) to 1 plus the number of total permutations (1000). Permutation test results provided in **Table S1.**

### Curation of Known DNA-Protein Binding Sites

Binding sites previously determined by global ChIP-seq studies (LexA^16^, mIHF^17^, HupB^18^, all others from ^6^), or manually curated from DNAse footprinting gels if crosslinking incompatible (IdeR^24^), were used to compare to IPOD-HR binding signal. All datasets used were obtained in *M. tuberculosis* H37Rv strain (accession either NC_000962.2 or NC_000962.3), and thus binding sites sequences were converted to *M. bovis* BCG coordinates (accession: AM408590.1). BLASTn ^43^ was used via command line with the following inputs:

blastn -db ./BlastDB/AM408590_1.fasta -query query.fasta -evalue 0.0001 -out ./BlastResults/results.txt

Hits were then filtered to only unique binding sites with maximum sequence homology (*i.e.*, results sorted by Bitscore, where the highest hit was considered the binding site; multiple maximum Bitscore sites were removed as non-unique). ∼97% of binding sites were mapped to unique sites in *M. bovis* BCG given the high genomic sequence homology with *M. tuberculosis,* representing a total of 159 regulators (sites listed in **Table S1**).

### Enrichment of Protein Binding Sites in EPODs, nEPODs, and Peak Calls

The curated DNA binding sites for the DNA-binding proteins were used as our ground-truth dataset to compare to regions called as peaks, EPODs, and nEPODs in the no stress IPOD-HR results. The bedtools intersect tool^42^ was used to determine overlaps (for EPODs and nEPODs, at least 50% of the binding site had to overlap). The recall (fraction of the curated binding sites that overlap with a called peak) for each protein was calculated. For enrichment of binding sites within EPODs, nEPODs, and peak calls, the log_2_ transformed ratio of the recall to the null (fraction of EPODs/nEPODs/peak call bps to the entire genome) was used to determine enrichment (positive values are enriched, negative values are unenriched). A 0.5 pseudocount was included to avoid undefined results for counts of 0. For peak calls, the FDR-corrected p-values for a binomial distribution for each regulator compared to the null (peak call genome coverage in the given condition) was calculated. Values for each protein for EPODs, nEPODs, and peak calls are provided in **Table S2**.

### Cross-species Comparisons

For comparisons between *M. bovis* BCG and *M. smegmatis* gene expression, homologous genes were defined by the BV-BRC Proteome Comparison tool utilizing BlastP^43,44^. Results of the proteome analysis (mapped to *M. tuberculosis* H37Rv genome) provided in **Table S5** and **Figure S7**. Sequence conservation between organisms (also mapped to the *M. tuberculosis* H37Rv genome) was determined using BV-BRC Genome Alignment tool using progressiveMauve^44,45^. Local alignment results are provided in **Table S5**. Statistical analysis for determining dependence of (n)EPOD regions on sequence conservation was calculated using contingency tables by Fisher Exact tests (p-value < 0.05). Regions were defined as conserved if >50% of region aligned between *M. bovis* BCG and *M. smegmatis*. Background regions were all regions not within EPODs or nEPODs in the given condition, excluding regions larger or smaller than the EPOD and nEPOD length distributions. Contingency table analysis results provided in **Table S5**.

### Gene Ontology Enrichment Analyses

For differential expression analysis as well as EPOD and nEPOD overlapping genes, Gene Ontology analysis was performed. First, gene annotation sets were derived using InterLabelGO+^46^ with a confidence score of 0.5 or greater. These results are provided in **Table S4**. Gene Ontology (GO) enrichment analysis was performed using iPAGE version 1.2a^47^. For differential expression, the Wald statistic output from the Sleuth analysis (b/se_b) was used as the normalized expression values, in which the continuous values were binned.

For the EPOD and nEPOD overlapping genes, genes were discretely categorized as 0 (not overlapping in either condition), 1 or 2 (overlapping in only one of the two conditions, or 3 (overlapping in both conditions) with the –exptype=discrete parameter. In all tested conditions, the redundancy filter was used to remove redundant GO terms. However, this filter can result in obscure GO terms with masking of more common/abundant terms and thus the summary terms with their removed redundant terms are provided in **Table S4**.

## SUPPLEMENTARY DATA

File S1. Combined supplementary information and supplemental figures.

Table S1. Protein occupancies. The (n)EPOD regions and permutation testing of GC content for *M. bovis* BCG and *M. smegmatis*. Peak calls, orphan peaks, and ChIP-seq mapped regulator binding sites in *M. bovis* BCG.

Table S2. Enrichment analysis results. Enrichment analysis of known protein binding sites within (n)EPODs, and peak calls, as well as the coordinates of dynamic (n)EPOD regions in *M. bovis* BCG.

Table S3. RNA-seq results. Transcript counts and differential expression results based on the assembled reference transcriptomes for *M. bovis* BCG and *M. smegmatis*. Inferred regulons for regulators based on binding site proximity to promoters and gene start sites.

Table S4. Gene Ontology results. The Gene Ontology annotations and results by iPAGE from the transcriptome and (n)EPOD analyses for *M. bovis* BCG and *M. smegmatis*.

Table S5. Organism comparison. Homologous genes between *M. bovis* BCG and *M. smegmatis*. Numerical results of the contingency table analysis of (n)EPOD conservation between organisms.

## AUTHOR CONTRIBUTIONS

Conceptualization, A.M.E, L.M.C and L.F; Methodology, A.M.E, J.W.S and L.F; Investigation, A.M.E, A.T, R.H, Q.L; Writing – Original Draft, A.M.E; Writing – Review & Editing, A.M.E, A.T, R.H, L.M.C and L.F; Funding Acquisition, L.M.C and L.F; Resources, L.M.C, and L.F.; Supervision, L.M.C and L.F

## ACKNOWLEDGMENTS

We would like to thank the Torreles Lab (Texas Biomedical Research Institute, San Antonio, TX) for sharing the *M. bovis* BCG Pasteur strain. The authors acknowledge the Texas Advanced Computing Center (TACC) at the University of Texas at Austin for providing HPC that supported the bioinformatic analyses of this work. Data analysis for this work also used the Advanced Cyberinfrastructure Coordination Ecosystem: Services & Support (ACCESS) program, which is supported by National Science Foundation (2138259, 2138286, 2138307, 2137603, and 2138296).

