## Supplementary material for "Conserved Heterochromatin-like Structures with Local Regulators Mediate the Iron Stress Response in Mycobacteria": EkdahlA_MycoIPODHR_SI_BioRxiv.pdf

**Supplementary Information -**

***Dynamic chromatin structures coordinating the oxidative stress response are captured by IPOD-HR***

Oxidative stress has previously been shown to induce large changes in chromatin structures in other bacteria[1–4]. Thus, we compared IPOD-HR gathered under oxidative stress compared to the no stress condition to ensure that IPOD-HR was able to capture stress-responsive chromatin structures in *M. bovis* BCG. To induce oxidative stress, cells were grown in parallel with the no stress samples and treated with 1mM tert-butyl hydroperoxide (TbHP) for 4 hours prior to sampling at mid logarithmic phase ($OD_{600}$ 0.6 – 1.0) as previously performed[5]. TbHP is an oxidizing reagent similar to, but more stable than, hydroperoxide. The greater stability of TbHP allows for longer oxidative stress application. We anticipated increased DNA condensation by proteins to protect the DNA from radical oxygen species as documented in prior literature[2].

As expected, oxidative stress resulted in genome-wide changes in gene expression and the DNA-protein landscape (**Figure S2A**). Notably, 1,031 transcripts encompassing 1,564 genes, out of a total 2,739 transcripts and 2,988 genes, were significantly differentially expressed (significance defined as $|\log_2 FC| > 1$ and p-adj <0.05), particularly including transcripts for induction of DNA damage and oxidative stress response genes during oxidative stress (**Figure S2B, Figure S3A**).

Alongside the sweeping changes in the transcriptome, numerous EPOD and nEPOD regions changed (**Figure S2A**). As noted in the main text, nEPODs are also referred to as mIHF-type nEPODs due to the high prevalence of mIHF binding sites in nEPOD regions. EPOD and mIHF-type nEPOD regions that formed during stress (188 and 295 EPODs and nEPODs, respectively) outnumbered those that dissipated during stress (120 and 67). Indeed, the average lengths of EPODs and nEPODs in oxidative stress (3.6 and 3.4kb, respectively) are significantly higher than in no stress (2.5 and 2.6kb). The observation of increased DNA protein occupancy during oxidative stress agrees with the expectation that oxidative stress would result in increased DNA condensation. Functionally, EPODs have been attributed to silencing the transcription of genes by preventing binding or stalling elongation of the RNAP from gene promoters[6–8]. This hypothesis is consistent with our results in *M. bovis* BCG during oxidative stress, where in **Figure S3B** we found that the expression of genes was relatively unchanged when genes were contained within EPODs in both tested conditions (green). However, we observed changes in gene expression in regions of dynamic EPODs: EPOD dissipation during oxidative stress resulted in positive $\log_2$FC values (grey) or increased transcription, and EPOD existing only during oxidative stress resulted in negative $\log_2$FC values (blue) or decreased transcription, consistent with condition-dependent gene silencing by EPODs when/where they are present. Indeed, the cellular pathways contained within EPODs (and their changes during stress) further support EPOD-dependent transcriptional regulation during oxidative stress (**Figure S2C**). We observed that enriched Gene Ontology terms within dynamic EPODs that dissipated upon stress (thus permitting transcription) included cellular response to DNA damage stimulus and oxidoreductase activity (redundant with the adhesion of symbiont to host term, orange arrows). In contrast, EPODs that formed upon stress (usually repressing transcription) included nucleotide metabolism (also redundant with the proton motive force-driven ATP synthesis term, black arrows). This pattern, however, did not hold for mIHF-type nEPODs, where genome-wide changes in constitutive versus dynamic nEPOD regions did not result in significant changes in expression. Furthermore, cellular pathways of genes within mIHF-type nEPODs were predominantly unchanged during stress, such as cell wall biogenesis and lipopolysaccharide metabolic processes (**Figure S2D**, grey arrows). One speculation is that perhaps the genes contained within mIHF-type nEPODs are not needed during the tested conditions. This tight interplay between local and global regulators for transcriptional regulation has been observed before in *E. coli*[9,10].

Surprisingly, upon exposure to TbHP, while the mIHF-type nEPOD GC content distribution remained significantly lower than the rest of the genome, the EPODs distribution reduced in GC content such that it was no longer significantly different from background (**Figure S3C**). The drop in GC content of the EPODs distribution upon oxidative stress suggests proteins contributing to EPOD formation selectively release from relatively high GC content regions of the DNA during oxidative stress (see **Figure 2F** and **Figure S4** on PE-PGRS loci), possibly reflecting differences in the main proteins forming those EPODs. A good example of a dynamic or stress-responsive EPOD with high GC content (69%) is the genomic region encoding two ATP-dependent DNA helicases (BCG_3226c and BCG_3227c), shown in **Figure S2E**. This operon was within an EPOD during the non-stress condition and not within an EPOD during oxidative stress. Consistent with the changes in protein occupancy, these helicase genes were significantly induced during oxidative stress ($log_2FC = 10.38$), also indicated by the increased RNAP binding throughout the operon. Notice the promoter showed similar RNAP binding in both conditions, thus the EPOD was likely restricting RNAP elongation. HupB has three mapped sites in this EPOD, suggesting HupB was a major contributor to the EPOD, and released from the DNA to enable transcription of the helicase genes to aid DNA repair.

Alternatively, the numerous cellular pathways enriched in differentially expressed transcripts (**Figure S2B**) that are not enriched in either EPODs or mIHF-type nEPODs (**Figure S2C-D**) suggest local transcription factor regulation of some essential pathways like translation (by the ribosome assembly and protein catabolic process terms) and cellular respiration (redundant with the generation of precursor metabolites and energy term). Our protein occupancy data did capture local transcription factor regulation particularly in open chromatin regions; we show an example in **Figure S2F** for the operon encoding the DNA damage-responsive alternative sigma factor, SigG[11,12]. This three gene operon (*sigG-BCG_0218c-BCG_0217C)* was induced during oxidative stress ($log_2FC = 3.22$), as is apparent by the appearance of RNAP binding at the promoter only during oxidative stress. Numerous regulators have ChIP-seq mapped binding sites near the operon (highlighted in the shaded blue regions). While we cannot determine the protein identities directly from the IPOD-HR dataset, we can infer their likely identities when in conjunction with the prior ChIP-seq binding sites. For example, two uncharacterized regulators, Rv0678 and Rv0691c, have binding sites just upstream of *sigG* in which protein binding occurred only during the no stress condition (shaded blue region) likely contributing to the repression of the operon when DNA damage is not present. Also notable of this operon is the nucleoid-associated protein NapM, which has two binding sites within the operon. It is postulated that these internal binding sites may compact this DNA region to also repress transcription during basal conditions alongside the local transcription factor regulation.

Overall, the protein occupancy dynamics during oxidative stress revealed both local transcription factor regulation and global chromatin structures coordinating the transcriptional response. Our results agree with prior literature findings of increased global DNA condensation during oxidative stress, perhaps to protect against DNA damage[1,2]. However, the EPODs appeared more stress-responsive, dissipating during oxidative stress to induce both DNA repair and oxidoreductase cellular pathways while the mIHF-type nEPODs appeared more constitutive, regulating processes such as cell wall composition that do not vary across these stresses (consistent with the iron stress observations in the main text).

Since high iron and oxidative stress are related through the Fenton Reaction[13], where excess intracellular iron ions catalyze the formation of radical oxygen species, we anticipated the iron-poor condition to contrast with the oxidative stress response pathways (*i.e.*, low iron levels would have reduced radical oxygen species). Surprisingly, contrary to our expectations, we found a positive correlation in log fold changes between 170 out of the 179 shared differentially expressed genes in both stress conditions (**Figure S3F**); that is, expression changes in these genes occurred in the same direction (*i.e.,* both up- or down-regulated) during oxidative and iron stress. Notably, 41 of the shared genes induced during both stresses are part of the DNA damage repair pathway (**Table S3**) including helicases and the SOS response regulator LexA and recombinase RecA, suggesting perhaps either an adapted anticipatory response[5,14] where iron-poor conditions induce a preparatory DNA repair response to expected oxidative stress, or an induction of endogenous radical oxygen species generation when iron levels are low, such as was found to occur in other species like *Anabaena*[15] and *C. crescentus*[16]. Furthermore, changes in the transcriptome were less extensive during response to iron stress as compared with oxidative stress (**Figure S3A** compared to **Figure S3D**, **Table S3**) with only 258 out of 2,739 transcripts differentially expressed, encoding for 405 genes (recall oxidative stress resulted in differential expression of 38% of the transcriptome). The abundance of transcripts encoded within dynamic EPODs and nEPODs was also less impacted during iron stress than during oxidative stress, where genes within EPODs were not significantly induced upon EPOD dissipation although the general trend remained (p-values of 0.13 and 0.19 for genes within stress-forming and stress-dissipating EPODs, respectively, **Figure S3E**).

***Considerations and limitations of IPOD-HR analysis in mycobacteria***

This study, for the first time, successfully applied the IPOD-HR experimental technique to *M. bovis* BCG to capture the global DNA protein occupancy at native conditions, without the need to overexpress or tag proteins. By leveraging ChIP-seq identified binding sites, IPOD-HR can be applied across differing growth conditions without the need to re-perform ChIP-seq experiments. However, a major limitation of IPOD-HR is the lack of protein identification.

It is important to acknowledge that our analysis relies on outside ChIP-seq datasets and how each study calls binding sites to propose protein identities. For example, the ChIP-seq identified binding sites' average widths are ~410 bp as opposed to the average 60 bp resolution of peak calls by IPOD-HR, due to the stronger digestion generally used during IPOD-HR. This size discrepancy made it challenging to assign individual IPOD-HR peak calls to a single ChIP-seq regulator binding site in this study. Alternatively, the EPOD region that covered the PE-PGRS3 and PE-PGRS4 operon in **Figure 1C** lacked mapped protein binding events to account for the entire occupied region. It is possible that broad HupB binding could contribute to the occupancy signal in the EPOD; NAPs tend not to appear as formal ChIP peaks and thus may not be defined as binding sites in the original study. Alternatively, there may be bound regulators that currently lack mapped ChIP-seq binding sites and thus cannot be identified within our dataset[6].

Given the high prevalence of negative IPOD signal in *M. bovis* BCG and *M. smegmatis*, it is likely that at least some DNA-binding proteins partition away from the extracted interphase (although given the presence of mIHF binding sites in more than 90% of nEPODs, mIHF may represent a rare but highly abundant example of this phenomenon). We have hypothesized that the interphase depletion of some proteins may be due to the protein chemistry in which the protein-DNA complexes preferentially reside in the aqueous layer instead of the interphase. Since peak calling was performed on the positive IPOD-HR signal, binding sites of proteins that partition away from the interphase were missed, likely accounting for some of the lower regulator binding site capture rates (**Figure S1B**, **Table S2**).

Comparing protein occupancies between *M. bovis* BCG and *M. smegmatis* can be challenging due to the significant differences between genomes, including the large genome reduction of *M. bovis* BCG as is common with pathogens, as well as the many gene and operon duplications, inversions, insertions, and deletions. While it is assumed protein occupancies and (n)EPOD thresholds are comparable between organisms, as the average protein occupancies of comparable (n)EPOD regions between organisms were relatively equal (**Figure S5F**), it is cautioned to interpret the data between organisms rather holistically (*i.e.,* large trends and patterns across operons and cellular pathways) instead of at the level of single protein occupancy values or peaks.

**Supplementary Table Descriptions -**

**Table S1. Protein occupancies.** The (n)EPOD regions and permutation testing of GC content for *M. bovis* BCG and *M. smegmatis*. Peak calls, orphan peaks, and ChIP-seq mapped regulator binding sites in *M. bovis* BCG.

**Table S2. Enrichment analysis results.** Enrichment analysis of known protein binding sites within (n)EPODs, and peak calls, as well as the coordinates of dynamic (n)EPOD regions in *M. bovis* BCG.

**Table S3. RNA-seq results.** Transcript counts and differential expression results based on the assembled reference transcriptomes for *M. bovis* BCG and *M. smegmatis*. Inferred regulons for regulators based on binding site proximity to promoters and gene start sites.

**Table S4. Gene Ontology results.** The Gene Ontology annotations and results by iPAGE from the transcriptome and (n)EPOD analyses for *M. bovis* BCG and *M. smegmatis*.

**Table S5. Organism comparison.** Homologous genes between *M. bovis* BCG and *M. smegmatis*. Numerical results of the contingency table analysis of (n)EPOD conservation between organisms.

### Supplementary Figures –

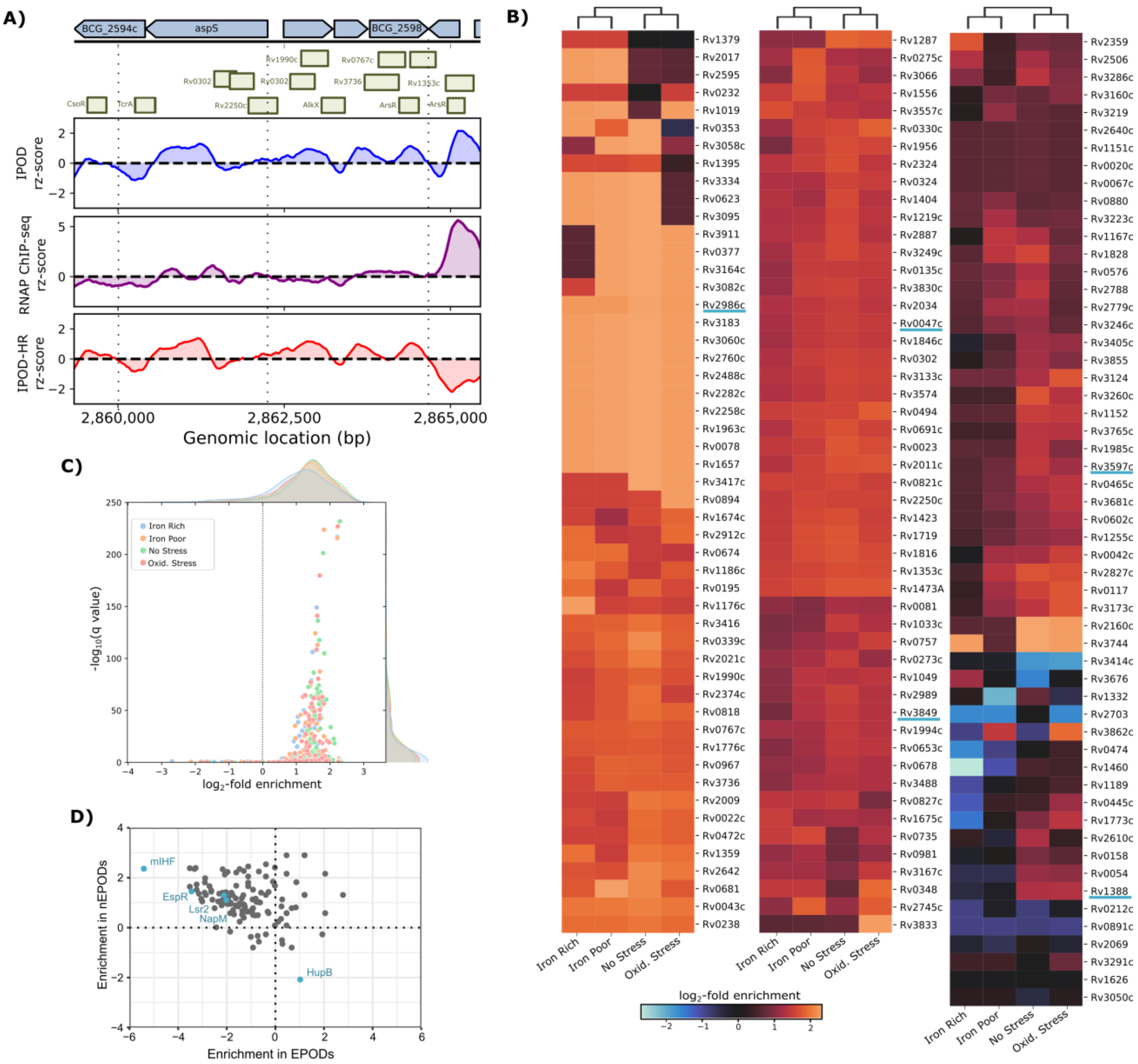


**Figure S1. Comparison of protein occupancy results to previously mapped protein binding sites. A)** Example of local regulatory logic captured by IPOD-HR, as illustrated with prior binding sites from outside ChIP-seq datasets (olive horizontal bars, sequence homology by BLASTn from *M. tuberculosis* datasets, annotated with *M. tuberculosis* gene names, **Table S1**) compared to the IPOD (blue), RNAP ChIP-Seq (purple), and IPOD-HR (red) results, all as 50 bp moving-averages. Select genes within depicted operon are annotated at top of panel. Note the IPOD occupancy peak (blue, top) is adjusted using the RNAP occupancy (purple, middle), resulting in no protein peak at a likely transcription start site (far right) in the final IPOD-HR occupancies (red, bottom). **B)** Clustered heat maps showing the $\log_2$-fold enrichment of overlaps between each transcription factor for which we were able to obtain data from prior ChIP-seq datasets (annotated by the *M. tuberculosis* gene name, conversions to *M. bovis* BCG names in **Table S1**), relative to the overall coverage of peaks on the genome under the corresponding condition. Key NAPs (HupB/Rv2986c, mIHF/Rv1388, Lsr2/Rv3597c, NapM/Rv0047c, EspR/Rv3849) are indicated with blue underlines. Enrichment data provided in **Table S2**. **C)** Comparison of peaks called from IPOD-HR occupancy signals for all tested conditions to known ChIP-seq mapped binding sites, plotted for enrichment values (x-axis; $log_2$-fold enrichment relative to the overall fraction of the genome covered by peaks) and statistical significance of the overlap (y-axis; based on a binomial test of the count of peaks overlapping known binding sites for each regulator, comparing the rate with the baseline rate of peak coverage; $-log_{10}$ FDR-corrected p-values are shown). **D)** Enrichment analysis of the regulator binding sites within EPODs (x-axis) and nEPODs (y-axis), calculated as the $log_2$(recall/null) for each regulator with 0.5 pseudocount to avoid log(0) errors. Positive values indicate a higher prevalence of binding sites within (n)EPODs compared to the rest of the genome. Known NAPs are highlighted in blue to emphasize the enrichment of regulator and NAP binding sites within nEPODs, compared to EPODs. Only HupB is found to be enriched in EPODs out of the 5 studied NAPs.

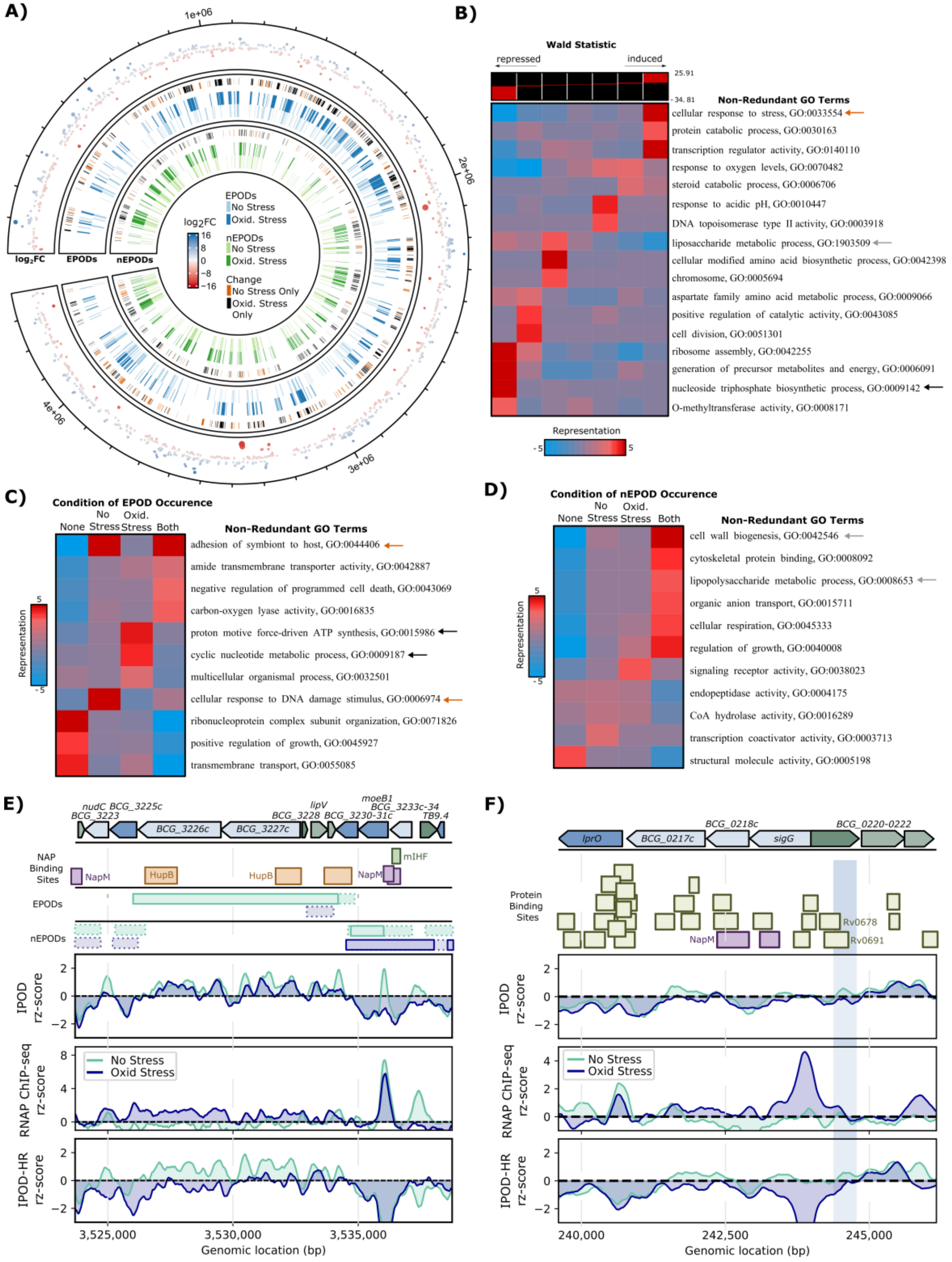


**Figure S2. Transcriptional response to oxidative stress is partially regulated by heterochromatin-like rearrangements. A)** Genome-wide depiction of the significant transcriptional changes (outer, p-adj < 0.05, blue is induced, red is repressed), EPODs (dark blue = oxidative stress, light blue = no stress), and nEPODs (dark green = oxidative stress, light green = no stress). The outer edge of the EPODs and nEPODs rings indicate regions >1kb that change between conditions (orange = (n)EPOD region only called in the no stress condition, black = region only called in the oxidative stress condition). **B)** Gene Ontology analysis of the RNA-seq differential expression results for oxidative stress. Noted by color coordinated arrows (with panels C-D) are cellular pathways discussed in **Supplementary Information** to be regulated by EPODs and nEPODs. **C-D)** Gene Ontology results of the genes encoded within (C) EPODs and (D) nEPODs during oxidative stress and no stress in *M. bovis* BCG. Columns indicate the conditions at which genes overlap with an (n)EPOD. GO term enrichment designated by color (red is overrepresented, blue is underrepresented). Orange arrows note cellular pathways transcriptionally induced by the dissipation of EPODs during oxidative stress, while black arrows note pathways repressed by the formation of EPODs during oxidative stress. **E)** Protein occupancy in no stress (green) and oxidative stress (blue) of the genomic region encoding two ATP-dependent helicases (BCG_3226c, BCG_3227c). Known binding sites of nucleoid-associated proteins (NAPs) depicted above the regions of strict (solid) and loose (dashed border) EPODs and nEPODs (see Methods for EPOD and nEPOD loose and strict threshold definitions). Notice the release of the EPOD, likely composed of HupB (orange) from no stress to oxidative stress coincides with an increase in RNA polymerase (RNAP) activity throughout the helicase genes. These genes are transcribed together with a measured $\log_2$FC of 10.38 during oxidative stress (p-adj = 2.8e-9). Note all genes surrounding the helicases, from *moeB1* through *nudC*, are likewise significantly induced (p-adj < 0.05, |log2fc| > 1). **F)** Protein occupancy in no stress (green) and oxidative stress (blue) of the genomic region encoding the DNA damage-responsive alternative sigma factor *sigG*. Note the numerous proteins that have mapped binding sites within and around the *sigG-BCG_0218-BCG_0217* operon (olive bars), including the NAP NapM (purple bars), many of which align with positive protein occupancy peaks in the IPOD-HR signal that occur only during the no stress condition. For example, as highlighted in the blue shaded region, two uncharacterized regulators, Rv0678 and Rv0691c, have binding sites just upstream of the likely transcription start site of the operon. Notably, oxidative stress results in high RNAP binding and transcription ($\log_2$fc = 3.22), as expected from the increased DNA damage, likely aided by the loss of transcriptional repressor binding of Rv0678 and/or Rv0691c

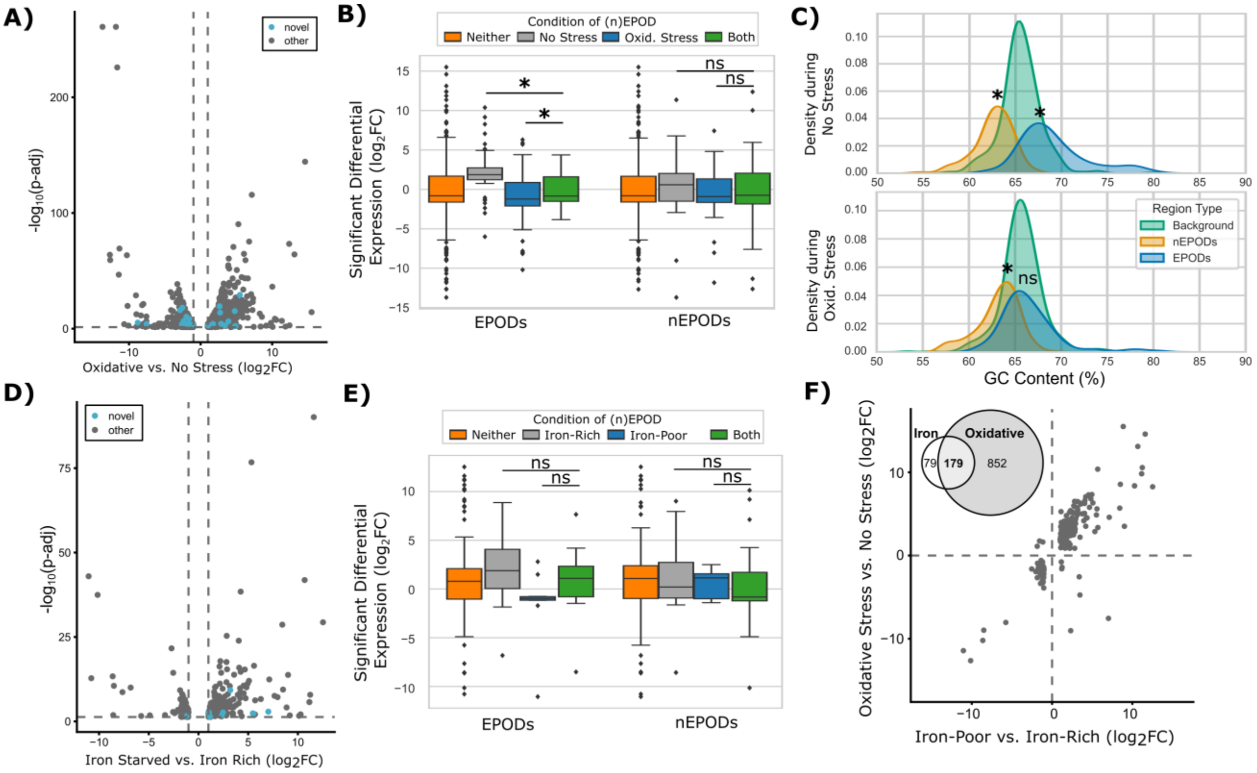


**Figure S3.** ***M. bovis*** **BCG transcriptional and protein occupancy responses to stress. A)** Volcano plots of significant differentially expressed transcripts ($|log_2FC| > 1$, p-adj < 0.05) during oxidative stress. Novel transcripts (*i.e.*, those newly identified in our transcriptome assemblies) are indicated in blue. **B)** The differential expression of genes (p-adj < 0.05) that are not contained in an EPOD (left) or nEPOD (right) in either condition (orange), only during no stress (grey), only during oxidative stress (blue), or in both conditions (green) conditions. Significance by two-tailed Welch's t-test is shown (comparisons with p-value < 0.05 with asterisk). Notice that EPOD occurrence significantly reduces transcription of genes, while nEPODs do not significantly alter expression in a consistent manner. **C)** Density plot of the GC content within EPODs (blue) and nEPODs (orange) compared to the rest of the genome (background, green) in no stress (top, duplicate of **Figure 1E**) compared to oxidative stress (bottom). Significance between EPODs and nEPODs to background was performed by permutation tests (p-val < 0.05). Notice EPODs, while significantly higher in GC content during non-stress conditions, are not significant during oxidative stress compared to background. This contrasts with the unchanging nEPOD region GC content, which stays significantly lower than background in both conditions. **D)** Volcano plots of significant differentially expressed transcripts ($|log_2FC| > 1$, p-adj < 0.05) during iron starvation. Novel transcripts are indicated in blue. **E)** The differential expression of genes (p-adj < 0.05) that are not contained in an EPOD (left) or nEPOD (right) in either condition (orange), only during iron-rich (grey), only during iron-poor (blue), or in both conditions (green) conditions. Significance by two-tail Welch's t-test is shown (comparisons with p-value < 0.05 with asterisk). Notice that while EPODs and nEPODs do not show statistical significance in regulating transcription in either condition, the pattern holds for EPODs for transcriptionally silencing transcription (p-values of 0.13 and 0.19, lack of significance likely due to smaller sample size with lower number of differentially expressed genes). **F)** Comparative scatter plot of the 179 shared differentially expressed transcripts ($|log_2FC| > 1$, p-adj < 0.05) during iron stress and oxidative stress. Venn diagram illustrates the number of transcripts differentially expressed in each condition independently and shared.

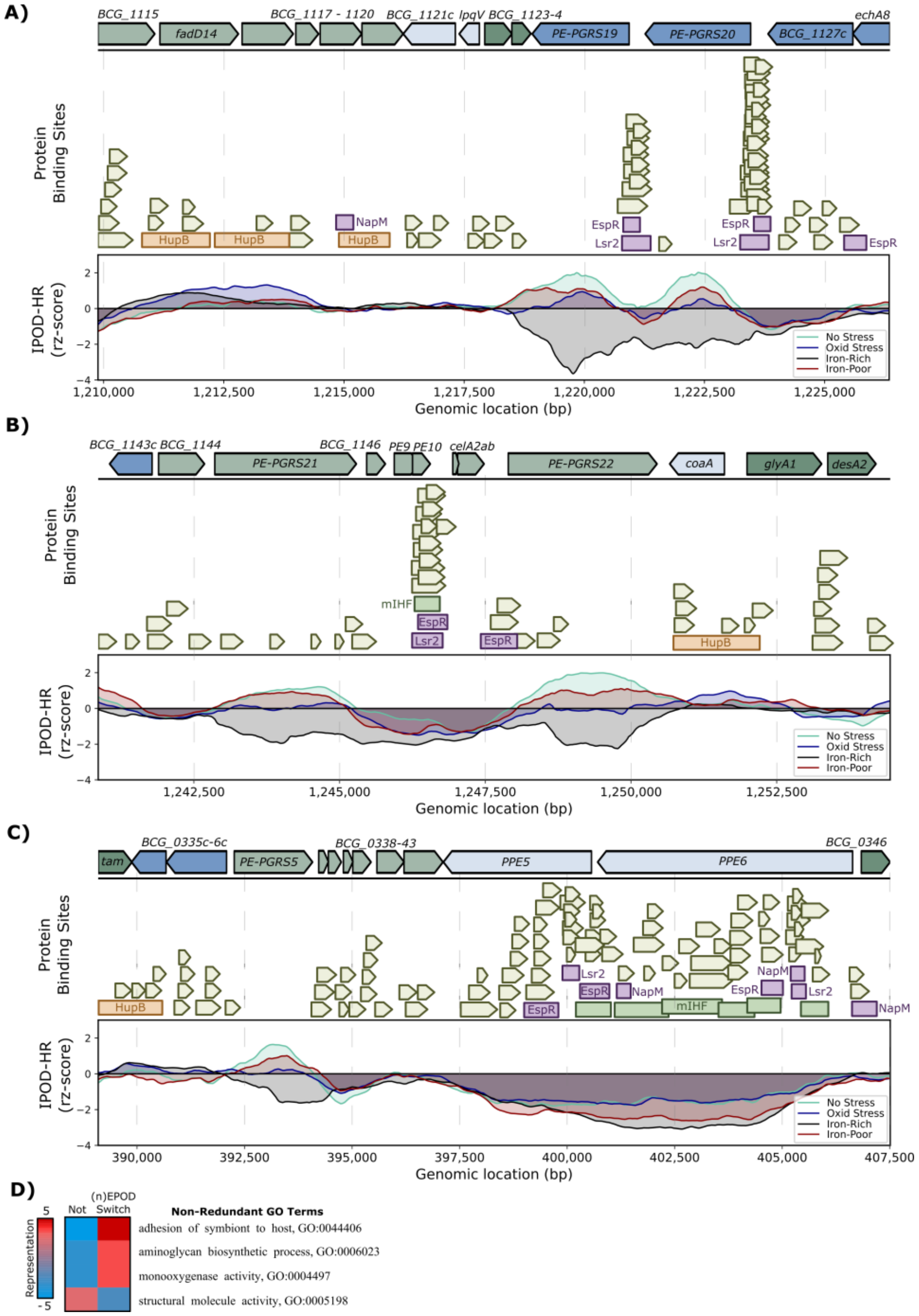


**Figure S4. Other EPOD and nEPOD switching examples at PE-PGRS loci in *M. bovis* BCG.** The IPOD-HR results with previously mapped protein binding sites around the operons encoding (A) the PE-PGRS19 and 20, (B) PE-PGRS21 and 22, and (C) PE-PGRS5 proteins. As observed in all three examples, highly bound protein occupancy over the entire PE-PGRS-encoded regions occurs during iron-poor and non-stress conditions (positive EPOD signals), as well as iron-rich (negative EPOD signals). Only in the oxidative stress condition do these regions lack protein occupancy. The dramatic changes in protein occupancy suggest dynamic protein binding events (from high to no protein occupancy) and changes in protein composition (EPOD to mIHF-type nEPOD signals). In all, fifteen PE-PGRS proteins (no. 5, 9, 10, 13, 14, 19-22, 27, 28, 33, 50, 52, 54) were encoded in regions that are within EPODs in one condition, and nEPODs during another.

**D)** Gene Ontology results of the genes contained within an EPOD or nEPOD and switches to the other during stress. The 376 and 249 genes that switch during iron and oxidative stress, respectively, alongside the PE-PGRS proteins (represented by the adhesion of symbiont to host pathway) include monooxygenase activity and aminoglycan biosynthetic processes pathways.

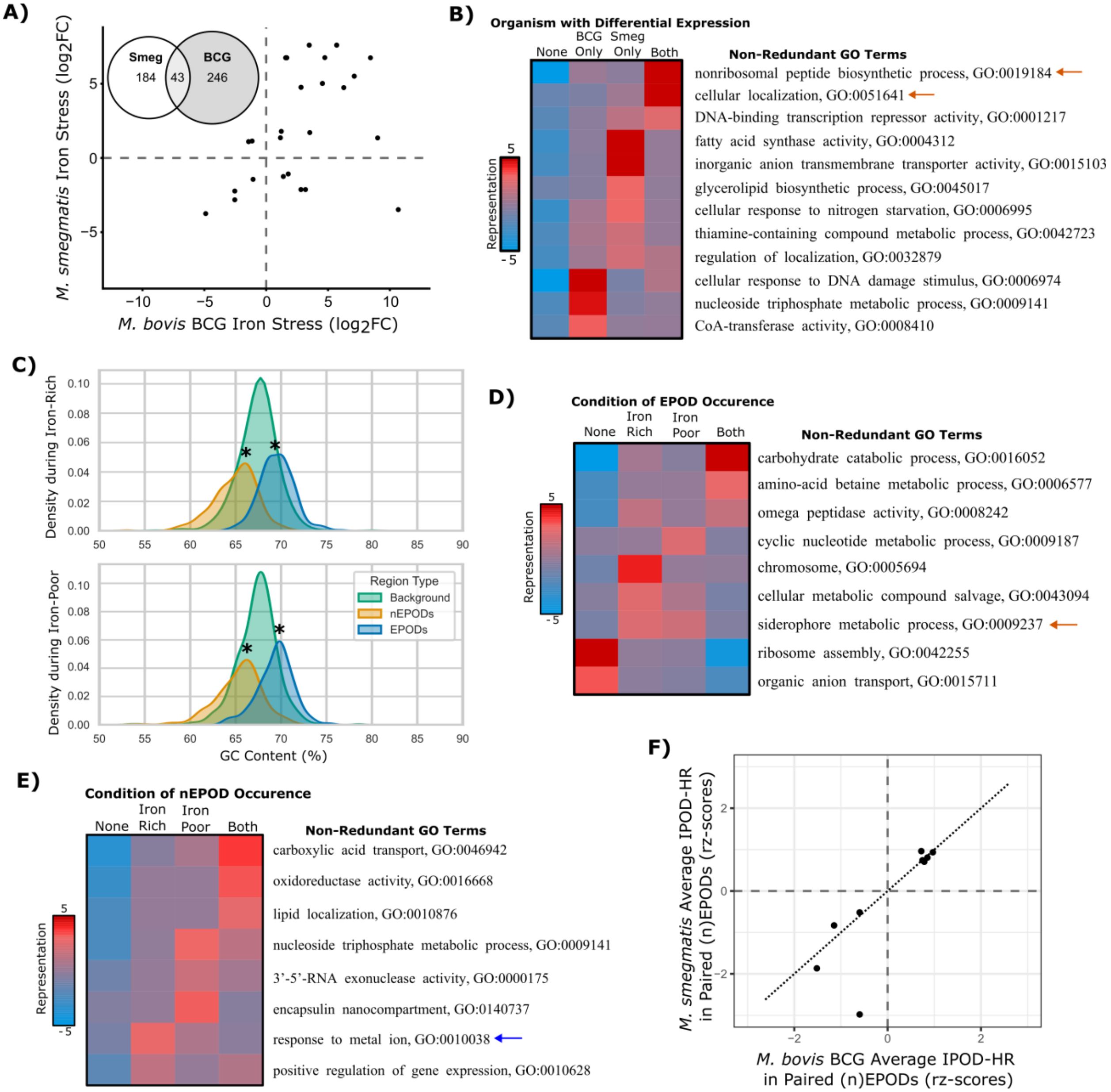


**Figure S5.** ***M. smegmatis* iron stress EPOD and nEPOD distributions as compared to *M. bovis* BCG. A)** Iron stress differential expression comparison between *M. bovis* BCG (x-axis) to *M. smegmatis* (y-axis) for homologous genes. Note the low overlap between organisms of only 43 genes, highlighted in the Venn Diagram. **B)** The Gene Ontology enrichment results of the significant differentially expressed genes during iron stress between *M. bovis* BCG and ***M. smegmatis*.** Representation of GO term shown by color (red: overrepresented, blue: underrepresented) for each bin (left: not differential in either organism, middle: differential in only one of the organisms as labeled, right: differential in both organism). **C)** Density plot of the GC content of EPODs, nEPODs, and background of *M. smegmatis* during iron-rich (left) and iron-poor (right) conditions. Note the nEPODs are significantly lower while EPODs are significantly higher in GC content compared to background (annotated by asterisk; $p<0.05$ by permutation test) in both conditions. **D-E)** Gene Ontology results by iPAGE [17] of the genes encoded within EPODs (D) and nEPODs (E) during iron-rich and poor conditions in *M. smegmatis*. Note the siderophore metabolic process pathway (orange arrow) shows some slight changes in EPODs, however to a much lesser extent compared to *M. bovis* BCG (only *mbtK* and *mbtN* were contained in EPODs during iron-rich conditions that dissipated to enable transcription in iron-poor conditions; this contrasts with the six mycobactin and carboxymycobactin genes in *M. bovis* BCG). As for nEPODs, or mIHF-type nEPODs, only slight iron-responsive regulatory behavior was supported by Gene Ontology similar to *M. bovis* BCG, including three genes of a cadmium regulatory operon (response to metal ion term, blue arrow) that were released from a mIHF-type nEPOD during iron stress. **F)** The average IPOD-HR occupancies (rz-scores) for comparable (n)EPODs between *M. bovis* BCG and *M. smegmatis*. Comparable (n)EPODs were defined as occurring across genomic sequences >50% conserved in both organisms. Notice the similar average protein occupancy (~1:1 relationship, dotted line) supporting the assumption of similar protein occupancy scores for comparing between the organisms.

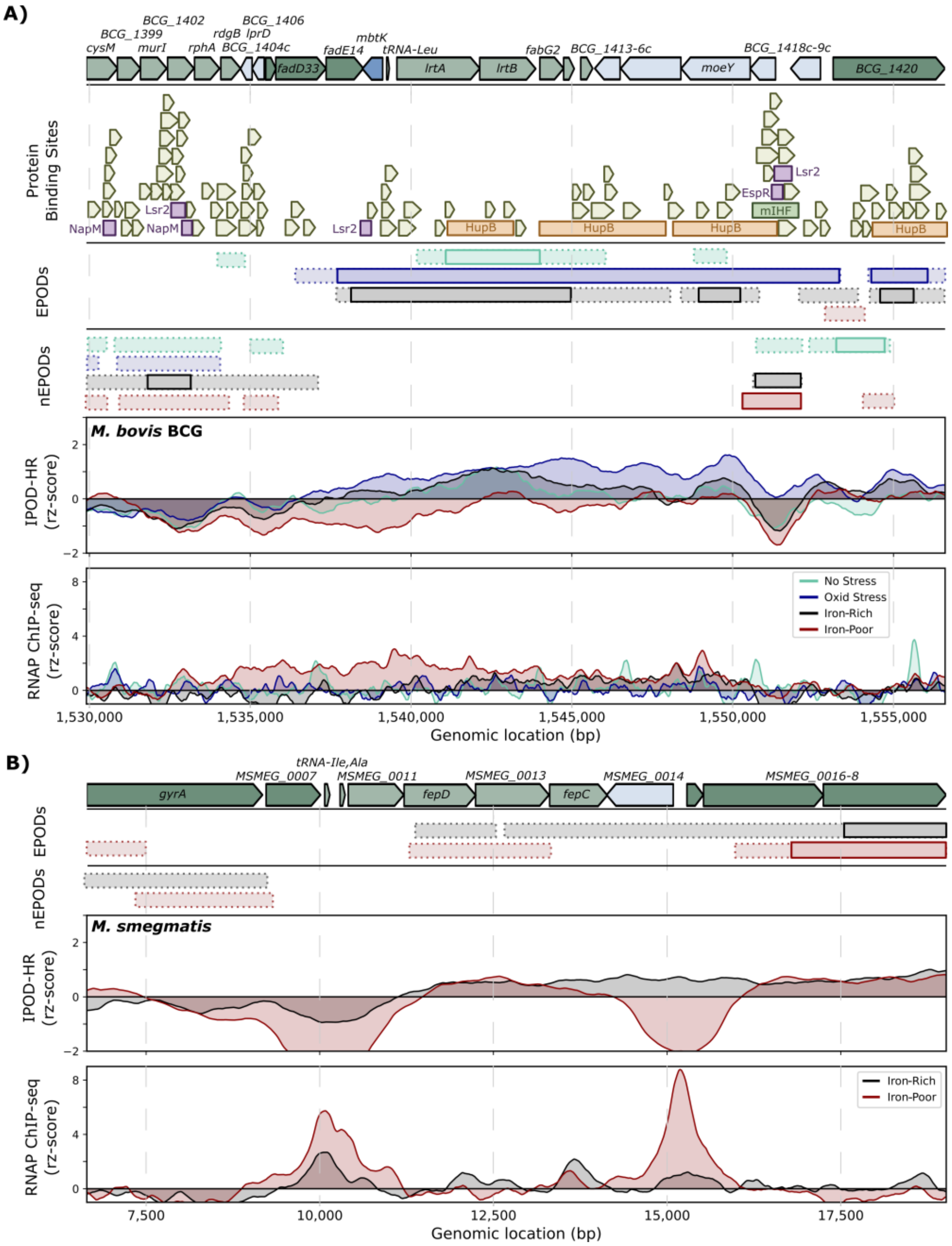


**Figure S6. Iron stress IPOD-HR results of siderophore synthesis operons that are not conserved between organisms. A)** The IPOD-HR results in the vicinity of the carboxymycobactin operon in *M. bovis* BCG for all tested conditions (no stress: green, oxidative stress: blue, iron-rich: black, and iron-poor: red). Note that *M. smegmatis* does not have a carboxymycobactin operon. Below the gene boundaries (blue and green arrows at top) are the previously mapped protein binding sites (olive green), including NAPs (purple, HupB: orange, mIHF: green), followed by the EPOD and then nEPOD regions (strict threshold with solid borders, loose threshold with dashed borders). The IPOD-HR scores (top panel) and RNAP ChIP-seq scores (bottom panel) emphasize the EPOD-directed transcriptional silencing of the carboxymycobactin operon, in which during iron starvation the protein occupancy is diminished to enable transcription (apparent by the increased RNAP activity heightened throughout operon). **B)** The IPOD-HR results of the exochelin operon (M*SMEG_0014* gene, also known as *fxbA*) in *M. smegmatis* for iron-rich (black) and iron-poor (red) conditions. Note *M. bovis* BCG does not have an exochelin operon. While there is a loose threshold EPOD apparent in iron-rich conditions (dashed) that is released in iron-poor conditions, the EPOD-dependent regulation is less striking than the *M. bovis* BCG siderophore examples. This operon is likely regulated in part by local transcription factors in addition to the weak EPODs.

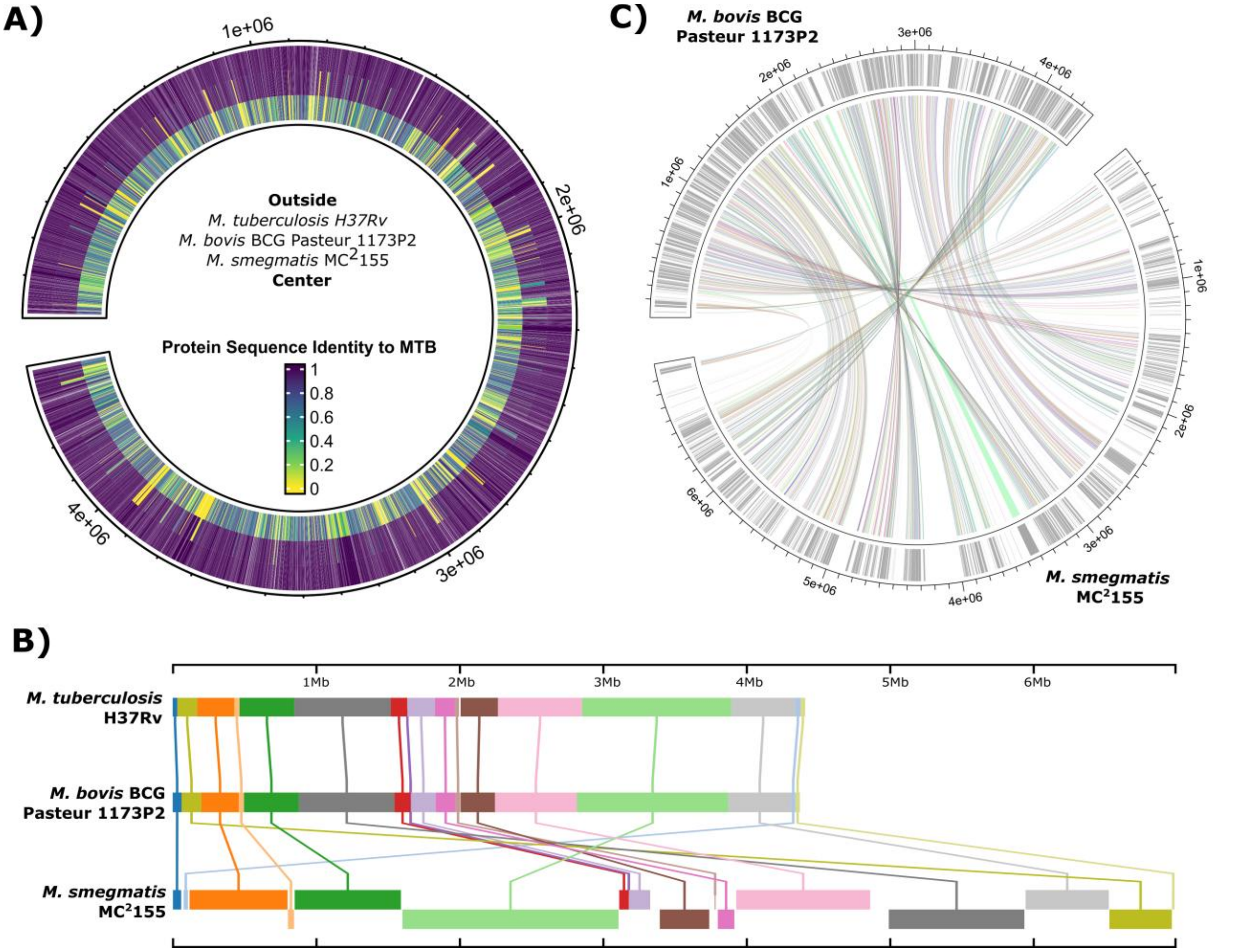


**Figure S7. BV-BRC Comparisons between organisms. A)** A circular diagram of the *M. tuberculosis* H37Rv genome (outside) comparing the protein sequence conservation of *M. smegmatis MC²155* (center) and *M. bovis* BCG Pasteur (middle) proteomes via the BV-BRC proteome comparison tool using BlastP[18]. Protein Sequence Identity (color) was calculated as the product of percent identity and percent coverage (data provided in **Table S5**). Note proteins with zero (yellow) Protein Sequence Identity indicate a lack of protein conservation to *M. tuberculosis*. **B)** An illustration of the local collinear blocks (LCBs) comparing the genome sequence similarity between the aforementioned Mycobacteria by the BV-BRC genome alignment tool using progressiveMauve[19] (seed length = 15). Note the much larger genome of the non-pathogen *M. smegmatis* as compared to the smaller *M. bovis* BCG (and *M. tuberculosis*) genomes (local alignments provided in **Table S5**). **C)** A more granular view of the conserved genomic regions between *M. bovis* BCG (top) and *M. smegmatis* (bottom) genomes. Grey regions (ring) illustrate regions that are conserved between the organisms, with randomly colored links mapping where the conserved regions are in each genome.

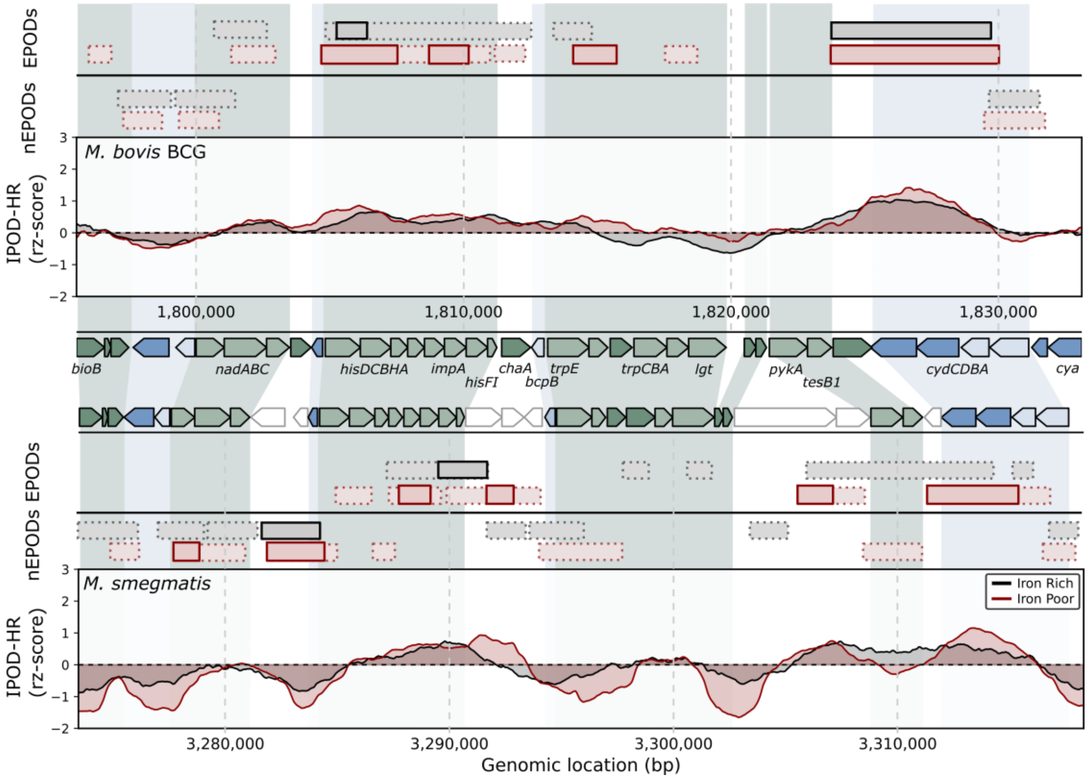


**Figure S8. Example of EPOD structure differences between *M. bovis* BCG and *M. smegmatis*: histidine and tryptophan biosynthesis operons.** IPOD-HR results around the histidine and tryptophan biosynthesis operons during iron-poor (red) and iron-rich (black) media conditions for *M. bovis* BCG (top) compared to *M. smegmatis* (bottom). Homologous genes shown by shading (green: forward strand, and blue: reverse strand) between organisms. Notice the differences in EPOD and nEPOD lengths and frequencies (horizontal boxes above the IPOD-HR results) between organisms, particularly in conserved (shaded) and non-conserved (white) regions. This example, while less striking than the cholesterol operon discussed in the main text, is more representative of average (n)EPOD lengths genome-wide for both organisms.

**Supplementary References -**
